# Integrative Transcriptomic and Functional Analysis Reveals Fatty Acyl Elongases Involved in Sex Pheromone Biosynthesis in Rice Leaffolder, *Cnaphalocrocis medinalis* (Lepidoptera: Pyraloidea)

**DOI:** 10.64898/2026.04.19.719439

**Authors:** Ling-Yue Chen, Xiao-Yang Lin, Ke-Xin Wang, Xiao Feng, Hong-Ting Tang, Shuang-Lin Dong, Ling-Ling Zheng, Yi-Han Xia

## Abstract

Elongases are essential enzymes in the biosynthesis of sex pheromones in many lepidopteran species. Together with desaturases, they determine the carbon skeletons of many pheromone precursors, thereby contributing to the production of species-specific chemical signals. However, to date, such fatty acyl elongase gene has not been functionally characterized. The rice leaffolder, *Cnaphalocrocis medinalis*, utilizes a blend of C18 monounsaturated aldehydes and alcohols as its sex pheromone, implying a critical elongation step from C16 precursors. In this study, we performed pheromone gland transcriptome analysis and identified 45 candidate biosynthetic genes. Functional assays in *Nicotiana benthamiana* showed that the Δ11 desaturase *Cmed070400* produces (Z)-11-hexadecenoic acid, which serves as the substrate for elongation. Multiple elongases catalyzed its conversion to (Z)-13-octadecenoic acid, with *Cmed092440* showing the highest activity. These findings provide the first experimental evidence for elongase-mediated formation of C18 pheromone precursors in *C. medinalis*. The identification of a minimal set of functionally active enzymes further enables reconstruction of this pathway in plant systems, offering a basis for sustainable production of pheromone precursors for pest management applications.

## Introduction

Moths rely on sex pheromones for mate communication, and synthetic pheromones are widely used in integrated pest management strategies such as monitoring, mass trapping, and mating disruption. Understanding the molecular basis of pheromone biosynthesis provides not only insights into chemical communication but also a foundation for the biotechnological production of species-specific pheromones [1–4].

Approximately 75% of the identified moth sex pheromones belong to Type I compounds, which are derived from palmitic and stearic acid through consecutive steps of fatty acyl desaturation, chain-shortening or chain-elongation, reduction and terminal functionalization [5]. Although pheromone biosynthetic pathways have been extensively investigated [6–10], functional characterization has largely focused on fatty acyl desaturases (FADs) [11–22], and fatty acyl reductases (FARs) [23–25], and a single example of chain-shortening enzyme (ACO) [11], whereas other enzymes in key steps, particularly fatty acyl elongases (ELOs) which responsible for the biosynthetic step of acyl-chain extension remain poorly understood [26–27].

Elongases are evolutionarily conserved enzymes found in animals, plants, and microorganisms, where they participate in the biosynthesis of saturated and unsaturated fatty acids, playing important roles in various biological processes. Historically, *ELO1* is the first identified membrane elongase in *Saccharomyces cerevisiae*, which is involved in the elongation of myristic acid to palmitic acid [28]. Thereafter, the *ELO2* and *ELO3* genes were identified by homology search based on the *S. cerevisiae* genome database [29]. In mammals, there are seven distinct ELO proteins that participate in biosynthesis of saturated, monounsaturated and polyunsaturated fatty acids [30–32]. To the best of our knowledge, the role of ELO genes in pheromone biosynthesis has not yet been reported in Lepidoptera. In other insect orders, however, several ELOs have been functionally characterized. For example, in *Drosophila*, *elo68α* is selectively expressed in the male reproductive system [33], while *eloF* is involved in long-chain hydrocarbon biosynthesis and courtship behavior in *D. melanogaster* [34]. In addition, *noa*, a homolog of mammalian *ELOVL6*, is required for viability and sperm development [35], and *bond* plays an essential role in *Drosophila* sex pheromone biosynthesis [36, 37]. Another example, *mElo*, contributes to desert adaptation in *D. mojavensis* [38]. Beyond *Drosophila*, heterologous expression of *TmELO1* from *Tenebrio molitor* L. in yeast enabled elongation of saturated fatty acids up to C24 [39], and elongation activities of *Elovl2/5*, *Elovl4* and *Elovl1/7* have been demonstrated in *Hediste diversicolor* [40]. In *Blattella germanica,* the female-enriched *BgElo12* is involved in maintaining sexually dimorphic hydrocarbons and female attractiveness [41]. These studies highlight the functional diversity of elongases in insects but also underscore the lack of experimental evidence for their role in pheromone biosynthesis in Lepidoptera.

The sex pheromone of the rice leaffolder *C. medinalis* (Guenée) (Lepidoptera: Pyraloidea) consists of a blend of C18 monounsaturated aldehydes and alcohols, which is a mixture of (Z)-13-octadecenal (Z13-18:Ald), (Z)-11-octadecenal (Z11-18:Ald), (Z)-11-octadecen-1-ol (Z11-18:OH) and (Z)-13-octadecen-1-ol (Z13-18:OH) at the ratio of 25: 3: 3: 3, suggesting that elongation of C16 precursors is a critical step in its biosynthesis [42, 43]. This species therefore provides a suitable system for investigating the role of elongases in pheromone production. In this study (Figure 1), we performed transcriptome sequencing of *C. medinalis* pheromone glands and identified 45 candidate genes involved in pheromone biosynthesis, including multiple elongase-like genes. Using gene expression profiling, phylogenetic analysis, and heterologous expression in *N. benthamiana*, we found that the Δ11 desaturase *Cmed070400* produces (Z)-11-hexadecenoic acid (Z11-16:Acid), which can be further elongated to (Z)-13-octadecenoic acid (Z13-18:Acid) by several elongases, with *Cmed092440* showing comparatively higher activity. These results highlight the role of elongases in determining pheromone precursor chain length within the biosynthetic pathway. Our study provides a molecular framework for future investigations of elongase function in moth pheromone biosynthesis and establishes a foundation for the bioproduction of the key precursor Z13-18:acid.

**Figure 1.**
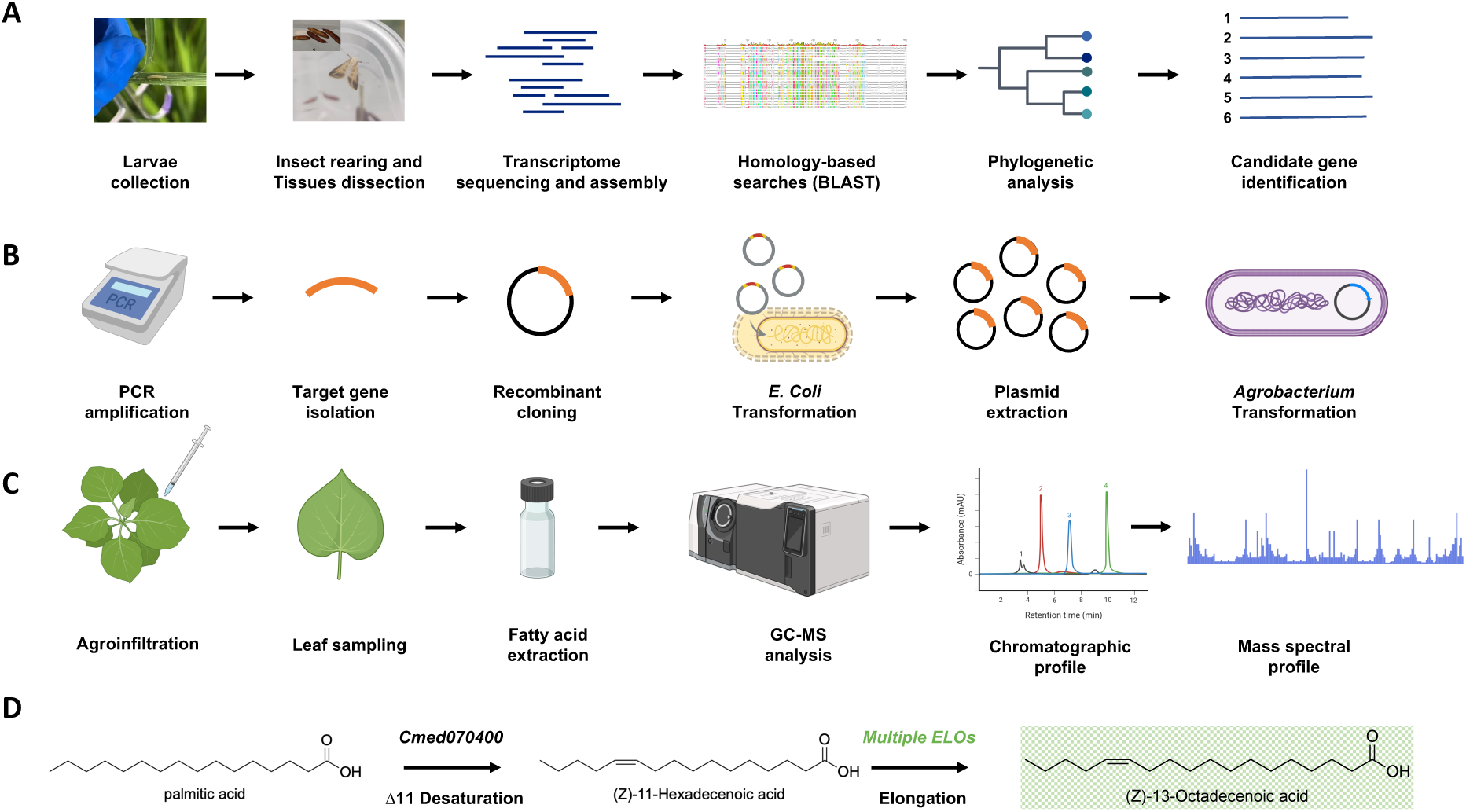
Experimental workflow and functional characterization of pheromone biosynthesis genes in *C. medinalis*. (A) Overview of transcriptome-based gene discovery. Larvae were collected from field, followed by insect rearing until adults for tissue dissection, RNA sequencing, and transcriptome assembly. Candidate genes were identified through homology-based searches (BLAST) and phylogenetic analysis. (B) Cloning and vector construction of candidate genes. Target genes were amplified, cloned into recombinant plasmids, and transformed into *Escherichia coli* for propagation, followed by plasmid extraction and transformation into *Agrobacterium tumefaciens*. (C) Functional characterization using a plant heterologous expression system. Recombinant *Agrobacterium* strains were used for leaf infiltration in *N. benthamiana*, followed by incubation with various gene combinations, leaf sampling, fatty acid extraction, and GC–MS analysis to determine product profiles. (D) Proposed biosynthetic pathway. The Δ11 desaturase Cmed070400 catalyzes the formation of (Z)-11-hexadecenoic acid, which is subsequently elongated by multiple elongases to produce (Z)-13-octadecenoic acid, the direct precursor of the major pheromone component in *C. medinalis*.

## Results

### Transcriptome overview of *C. medinalis*

In this study, 14 tissues of *C. medinalis* were collected, including heads, thoraxes, abdomens, legs, wings, and antennae from both sexes, as well as female pheromone glands (PG) and male hair-pencils (HP). Illumina sequencing generated a total of 654 million raw reads, of which over 96% were retained after quality filtering, yielding approximately 635 million high-quality clean reads (Fig. 2a, Table 1). These reads were de novo assembled into 86,256 unigenes and 225,513 transcripts (Fig. 2c). The sequencing data exhibited high quality, with an average error rate of 0.01 and Q20 and Q30 values of 98.81% and 96.63%, respectively, along with a GC content of 47.37% (Table 1). Additionally, mapping of clean reads to the reference genome resulted in an average alignment rate of 75.04% (consistent across replicates) (Fig. 2b). Furthermore, functional annotation against multiple databases showed that 56.10% of unigenes matched sequences in the NT database, followed by NR (34.72%) and PFAM and GO (both 23.72%) (Fig. 2d). Nearly half of the unigenes (49.40%) exhibited highest homology to lepidopteran species, predominantly *Ostrinia furnacalis* (35.70%) (Fig. 2f). The distribution of E-values further confirmed the robustness of sequence annotation (Fig. 2e).

**Figure 2.**
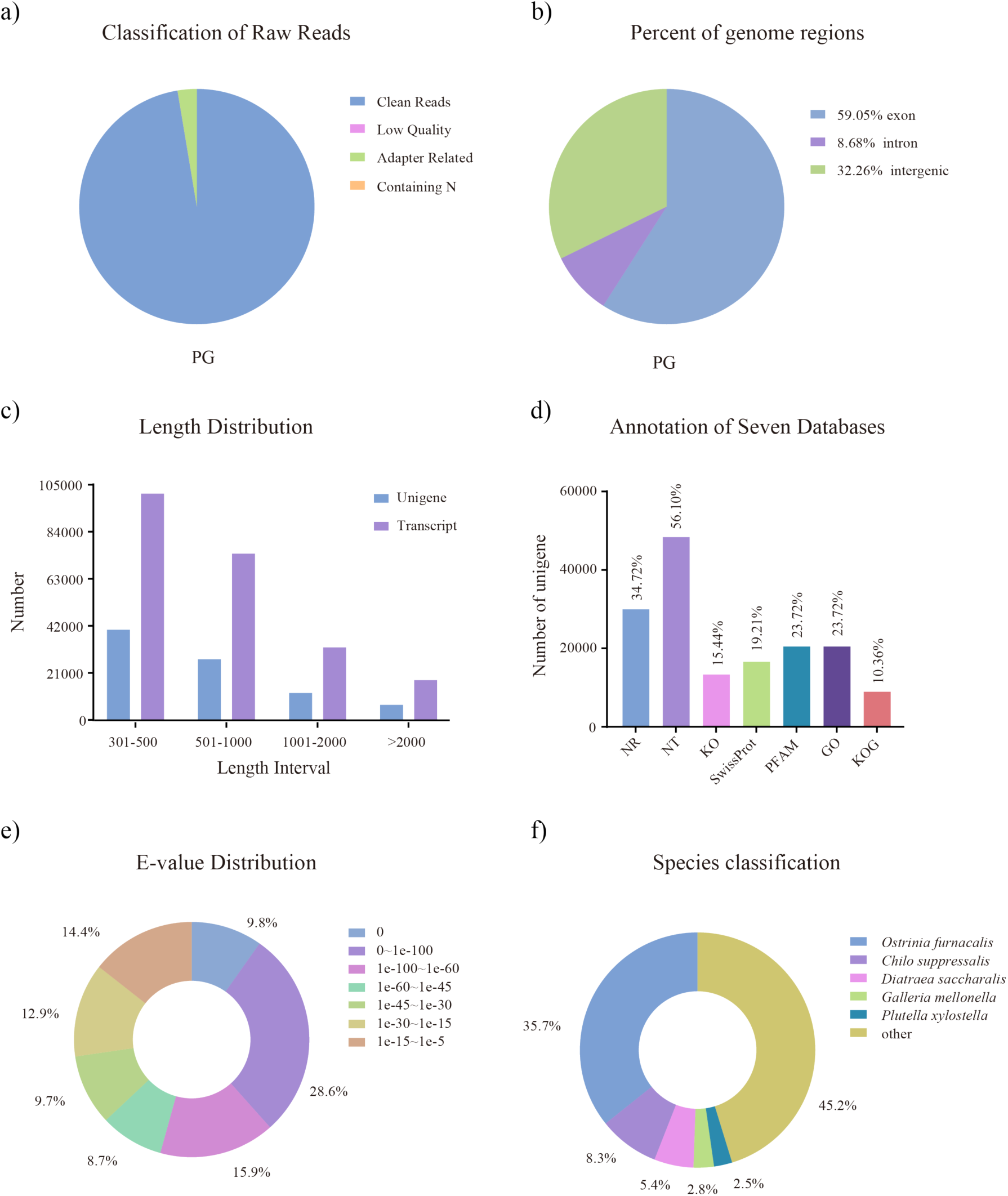
Transcriptome overview of *C. medinalis*. a) Composition of raw sequencing data from pheromone glands sample before filtering. The chart shows the relative proportions of clean reads (blue), low-quality reads (pink), adapter-containing reads (green), and reads with N bases (orange). The proportions of low-quality reads (pink) and reads with N bases (orange) are too low to be clearly visible in the chart. b) Genomic mapping of sequencing reads. The chart shows the relative proportions of reads aligned to exons (blue), introns (purple), and intergenic regions (green). c) Length distribution of transcripts and unigenes. The x-axis represents the length intervals, and the y-axis represents the number of sequences. d) Functional annotation of unigenes across seven databases. e) E value distribution of unigenes against the NCBI NR protein database. f) Species distribution of unigene homology, with the highest similarity to *Ostrinia furnacalis*.

**Table 1.** Summary statistics of transcriptome sequencing data from *C. medinalis* tissues.

| Sample | Raw reads number | Raw data bases(G) | Clean reads number | Clean data bases(G) | Error rate | Clean data Q20(%) | Clean data Q30(%) | Clean data GC(%) | Accession number (GSA) |
| --- | --- | --- | --- | --- | --- | --- | --- | --- | --- |
| F_A | 47824952 | 7.17 | 46749236 | 7.01 | 0.01 | 98.84 | 96.78 | 44.45 | CRA034037 |
| F_Ab | 43188016 | 6.48 | 41252388 | 6.19 | 0.01 | 98.77 | 96.52 | 48.19 | CRA034034 |
| F_H | 50349898 | 7.55 | 49624620 | 7.44 | 0.01 | 99.18 | 97.54 | 45.38 | CRA034032 |
| F_L | 48493142 | 7.27 | 47225522 | 7.08 | 0.01 | 98.84 | 96.71 | 48.87 | CRA034035 |
| F_PG | 41331526 | 6.2 | 40243152 | 6.04 | 0.01 | 98.77 | 96.47 | 48.06 | CRA034038 |
| F_T | 44561014 | 6.68 | 42820458 | 6.42 | 0.01 | 98.8 | 96.53 | 45.94 | CRA034033 |
| F_W | 50450036 | 7.57 | 49233506 | 7.39 | 0.01 | 98.64 | 96.17 | 46.18 | CRA034036 |
| M_A | 47383618 | 7.11 | 46019206 | 6.9 | 0.01 | 98.83 | 96.73 | 44.71 | CRA034044 |
| M_Ab | 49151672 | 7.37 | 47492744 | 7.12 | 0.01 | 98.6 | 96.16 | 49.28 | CRA034041 |
| M_H | 49435078 | 7.42 | 48448786 | 7.27 | 0.01 | 99.13 | 97.42 | 45.71 | CRA034491 |
| M_HP | 43320168 | 6.5 | 42030862 | 6.3 | 0.01 | 98.62 | 96.14 | 48.91 | CRA034045 |
| M_L | 44621200 | 6.69 | 43189864 | 6.48 | 0.01 | 98.76 | 96.52 | 50.06 | CRA034042 |
| M_T | 46283546 | 6.94 | 44470890 | 6.67 | 0.01 | 98.77 | 96.48 | 49.88 | CRA034040 |
| M_W | 47810176 | 7.17 | 46518478 | 6.98 | 0.01 | 98.8 | 96.65 | 47.54 | CRA034043 |

Tissues are abbreviated as follows: F, female; M, male; H, head; T, thorax; Ab, abdomen; L, legs; W, wing; A, antennae; HP, hair-pencil, PG, pheromone gland. Raw reads number: the total number of raw sequencing reads generated for each sample. Raw data bases (G): total raw sequencing data generated for each sample in gigabases (G). Clean reads number: the total number of high-quality reads after filtering. Clean data bases (G): high-quality sequencing data after filtering in gigabases (G). Error rate refers to the average base-calling error per base. Clean data Q20 (%) and Q30 (%) indicate the percentage of bases with quality score $\geq 20$ and $\geq 30$ in clean data, respectively. Clean data GC (%): GC content of the clean data. The sequencing data for each tissue have been deposited in the Genome Sequence Archive (GSA) under corresponding accession number.

### Identification of candidate pheromone biosynthetic genes

Through homology-based searches, a total of 45 genes putatively involved in sex pheromone biosynthesis were identified, including gene candidates encoding 1 acetyl-CoA carboxylase (ACC), 4 fatty acid synthases (FAS), 12 FADs, 6 ACOs, 15 ELOs, 3 FARs, and 4 alcohol dehydrogenases (ADH) (Table 2). These gene families collectively represent key steps in the pheromone biosynthetic pathway. Upstream enzymes such as ACC and FAS are involved in fatty acid precursor synthesis, while desaturases introduce double bonds at specific positions. Notably, the identified FADs include both Δ9 and Δ11 subtypes, which are typically associated with pheromone precursor modification in moths. Elongases constitute a relatively large gene family (n = 15) and are likely responsible for extending fatty acyl chains from C₁₆ to C₁₈, a critical step for the production of *C. medinalis* pheromone components. Downstream, FARs and ADHs are expected to catalyze the conversion of fatty acyl intermediates into alcohols and aldehydes. Taken together, these results provide a comprehensive set of candidate genes spanning the entire pheromone biosynthetic pathway and highlight elongases as a prominent group potentially involved in determining pheromone precursor chain length (Table 2).

**Table 2.** Putative pheromone biosynthesis genes in the pheromone gland of *C. medinalis*.

| Best Blastx Match |  |  |  |  |  |  |
| --- | --- | --- | --- | --- | --- | --- |
| Gene ID /<br>Name | ORF<br>(bp) | Name | Acc.number | Species | E value | Identity(%) |
| **acetyl-CoA carboxylase(ACC)** |  |  |  |  |  |  |
| Cmed021440 | 1932 | acetyl-CoA carboxylase isoform X1 | XP_028156232.1 | *Ostrinia nubilalis* | 0.0 | 92.50 |
| **fatty acid synthase (FAS)** |  |  |  |  |  |  |
| Cmed153650 | 6777 | fatty acid synthase-like isoform X1 | XP_063833375.1 | *O. nubilalis* | 0.0 | 51.13 |
| Cmed097470 | 6184 | fatty acid synthase-like | XP_075991465.1 | *Anticarsia gemmatalis* | 0.0 | 47.89 |
| Cmed019060 | 2878 | fatty acid synthase-like isoform X2 | XP_063832938.1 | *O. nubilalis* | 0.0 | 83.39 |
| Cmed019050 | 2879 | fatty acid synthase-like isoform X1 | XP_063832937.1 | *O. nubilalis* | 0.0 | 94.64 |
| **Fatty Acyl Desaturase(DES)** |  |  |  |  |  |  |
| Cmed009890 | 999 | acyl-CoA Delta(11) desaturase-like | XP_028166624.1 | *O. furnacalis* | 0.0 | 88.55 |
| Cmed022650 | 1140 | acyl-CoA desaturase-like | XP_028162659.1 | *O. furnacalis* | 1e-144 | 54.24 |
| Cmed054120 | 1077 | acyl-CoA delta(11) desaturase 3 | UEN71133.1 | *Glyphodes pyloalis* | 0.0 | 93.58 |
| Cmed070400 | 1017 | acyl-CoA Delta(11) desaturase | XP_030022752.1 | *Manduca sexta* | 6e-160 | 62.80 |
| Cmed077910 | 903 | acyl-CoA Delta-9 desaturase-like | XP_063827537.1 | *O. nubilalis* | 6e-150 | 75.53 |
| Cmed116530 | 1113 | acyl-CoA Delta(11) desaturase-like; | XP_028172985.1 | *O. furnacalis* | 8e-165 | 71.63 |
| Cmed116590 | 1128 | acyl-CoA delta(11) desaturase 2 | UEN71132.1 | *G. pyloalis* | 0.0 | 82.14 |
| Cmed116600 | 1125 | (11Z)-hexadec-11-enoyl-CoA conjugase-like | XP_028172977.1 | *O. furnacalis* | 0.0 | 82.57 |
| Cmed138850 | 909 | acyl-CoA Delta(11) desaturase isoform X1 | XP_028176121.1 | *O. furnacalis* | 0.0 | 85.52 |
| Cmed034480 | 576 | acyl-CoA Delta(11) desaturase | XP_052745633.1 | *Bicyclus anynana* | 8e-46 | 68.52 |
| Cmed036160 | 804 | acyl-CoA desaturase-like | XP_028162659.1 | *O. furnacalis* | 7e-99 | 55.82 |
| Cmed116640 | 720 | acyl-CoA Delta-9 desaturase-like | XP_063822435.1 | *O. nubilalis* | 2e-133 | 74.90 |
| **Elongase (ELO)** |  |  |  |  |  |  |
| Cmed092430 | 987 | very long chain fatty acid elongase AAEL008004 isoform | XP_063890227.1 | *Helicoverpa armigera* | 0.0 | 91.37 |
| Cmed092440 | 933 | elongation of very long chain fatty acids protein | XP_047020111.1 | *H. zea* | 0.0 | 86.62 |
| Cmed092450 | 951 | very long chain fatty acid elongase AAEL008004 | XP_063820949.1 | *O. nubilalis* | 0.0 | 87.34 |
| Cmed153350 | 921 | elongation of very long chain fatty acids protein | XP_049865794.1 | *Pectinophora* | 2e-175 | 87.93 |
| Cmed153360 | 1296 | elongation of very long chain fatty acids protein | XP_028162527.1 | *O. furnacalis* | 8e-73 | 60.77 |
| Cmed153370 | 918 | elongation of very long chain fatty acids protein 4-like | XP_028162528.1 | *O. furnacalis* | 2e-137 | 65.73 |
| Cmed063180 | 810 | very long chain fatty acid elongase AAEL008004-like | XP_063833189.1 | *O. nubilalis* | 9e-167 | 94.42 |
| Cmed097790 | 666 | elongation of very long chain fatty acids protein 4-like | XP_028178350.1 | *O. furnacalis* | 4e-132 | 83.71 |
| Cmed140080 | 675 | elongation of very long chain fatty acids protein 4-like | NP_001299254.1 | *Papilio xuthus* | 2e-125 | 80.44 |
| Cmed153310 | 512 | elongation of very long chain fatty acids protein 7-like | XP_028162549.1 | *O. furnacalis* | 2e-94 | 80.34 |
| Cmed153340 | 880 | elongation of very long chain fatty acids protein 7-like | XP_028162542.1 | *O. furnacalis* | 2e-138 | 76.19 |
| Cmed153380 | 1770 | elongation of very long chain fatty acids protein 7-like | XP_047020502.1 | *H. zea* | 7e-175 | 77.27 |
| Cmed153390 | 714 | very long chain fatty acid elongase 4-like | XP_063839360.1 | *O. nubilalis* | 1e-119 | 61.81 |
| Cmed153400 | 684 | elongation of very long chain fatty acids protein | XP_030031441.2 | *M. sexta* | 2e-103 | 88.41 |
| Cmed039690 | 487 | elongation of very long chain fatty acids protein 6-like | XP_028173845.1 | *O. furnacalis* | 5e-110 | 96.88 |
| **acyl-coenzyme A oxidase (ACO)** |  |  |  |  |  |  |
| Cmed078630 | 2532 | acyl-CoA oxidase, partial | QZC92112.1 | *Dioryctria abietella* | 0.0 | 54.88 |
| Cmed078620 | 5610 | Acyl-coenzyme A oxidase, partial | KOB75219.1 | *Operophtera brumata* | 0.0 | 52.41 |
| Cmed078610 | 1653 | acyl-coenzyme A oxidase 1-like | XP_072946882.1 | *Epargyreus clarus* | 0.0 | 56.48 |
| Cmed078590 | 1971 | acyl-coenzyme A oxidase 1-like | XP_072946902.1 | *E. clarus* | 0.0 | 81.02 |
| Cmed010500 | 2496 | Peroxisomal acyl-coenzyme A oxidase 3 | KPJ19117.1 | *Papilio machaon* | 0.0 | 46.43 |
| Cmed010350 | 2052 | peroxisomal acyl-coenzyme A oxidase 3 | XP_028162586.1 | *O. furnacalis* | 0.0 | 85.78 |
| **fatty acyl reductase (FAR)** |  |  |  |  |  |  |
| Cmed135510 | 1544 | fatty acyl reductase 1, partial | UEV87807.1 | *Maruca vitrata* | 6e-173 | 82.27 |
| Cmed134040 | 1248 | fatty-acyl reductase | AGG19592.1 | *O. latipennis* | 1e-110 | 43.50 |
| Cmed075430 | 1569 | fatty acyl reductase 2 | UEV87808.1 | *M. vitrata* | 0.0 | 89.94 |
| **alcohol dehydrogenase (ADH)** |  |  |  |  |  |  |
| Cmed055150 | 1560 | type III alcohol dehydrogenase | XP_075970313.1 | *A. gemmatalis* | 0.0 | 79.09 |
| Cmed055160 | 1128 | type III alcohol dehydrogenase | XP_073957504.1 | *Choristoneura* | 0.0 | 62.99 |
| Cmed065450 | 1032 | zinc-type alcohol dehydrogenase-like protein SERP1785 | XP_034830422.1 | *Maniola hyperantus* | 0.0 | 72.67 |
| Cmed087870 | 1131 | alcohol dehydrogenase class-3 | XP_026762464.2 | *Galleria mellonella* | 0.0 | 91.22 |

For each gene, the open reading frame (ORF) length (bp) and the best match from a BLASTx hit against the NCBI database are listed.

The expression profiles of these genes in the pheromone gland are presented in Figures 3 and S2, showing consistent patterns across independent transcriptomic datasets. Notably, several FAD and ELO genes exhibited relatively high expression levels, with *Cmed070400* showing the highest abundance, consistent with their roles in fatty acyl desaturation and chain elongation. In contrast, genes associated with upstream and downstream steps generally showed lower or more variable expression levels. To validate the reliability of transcriptome-based expression estimates, several FAD and ELO genes were randomly selected for quantitative PCR (qPCR) analysis. The relative expression patterns obtained by qPCR were largely consistent with the FPKM values derived from RNA-seq data (Fig. 4), with highly expressed genes in the pheromone gland showing correspondingly elevated transcript levels. This concordance between qPCR and transcriptome data supports the reliability of the RNA-seq–based expression profiles for subsequent functional analysis.

**Figure 3.**
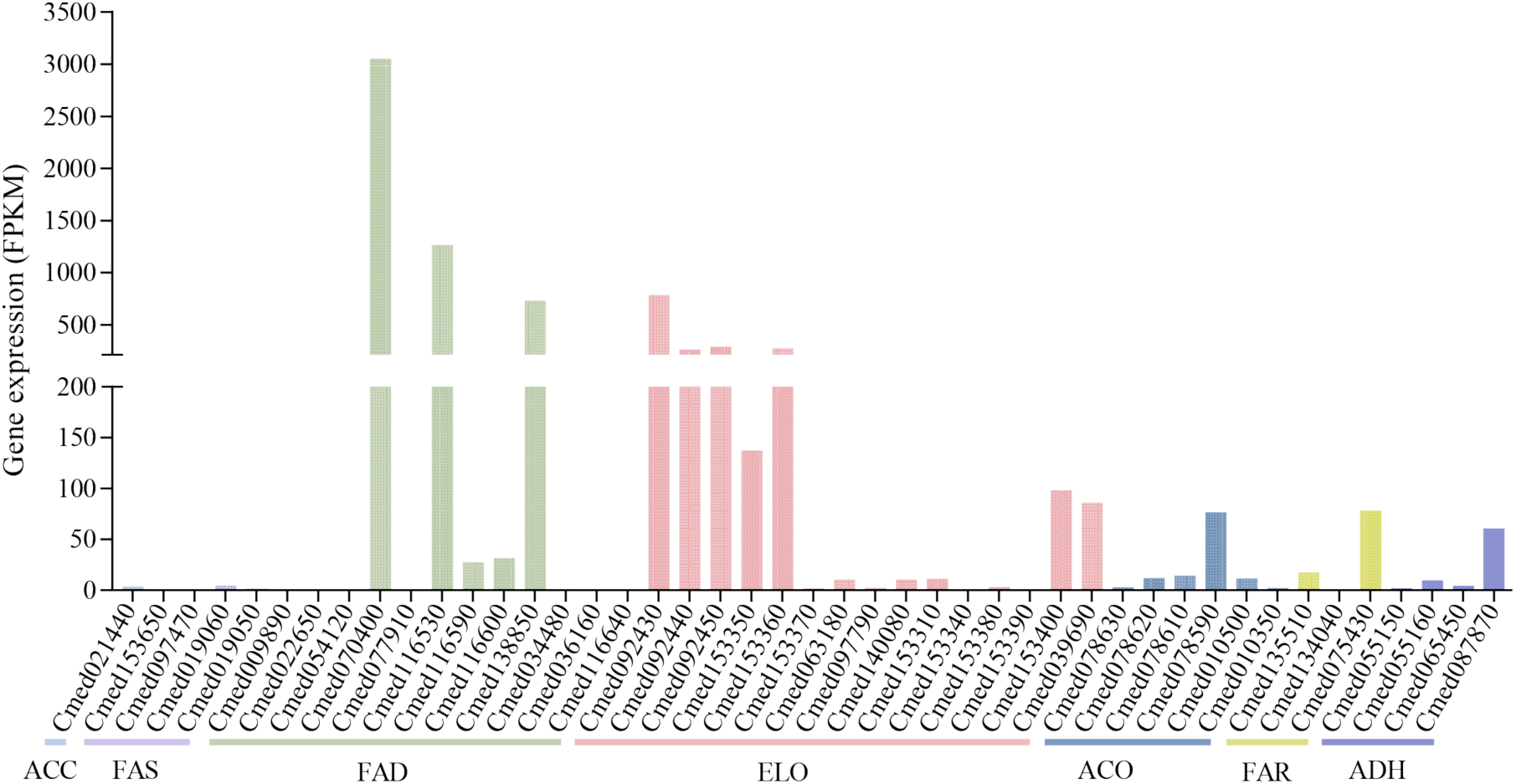
Expression levels of putative pheromone biosynthesis genes in the pheromone gland of *C. medinalis*. Gene expression levels are shown as Fragments Per Kilobase of transcript per Million mapped reads (FPKM).

**Figure 4.**
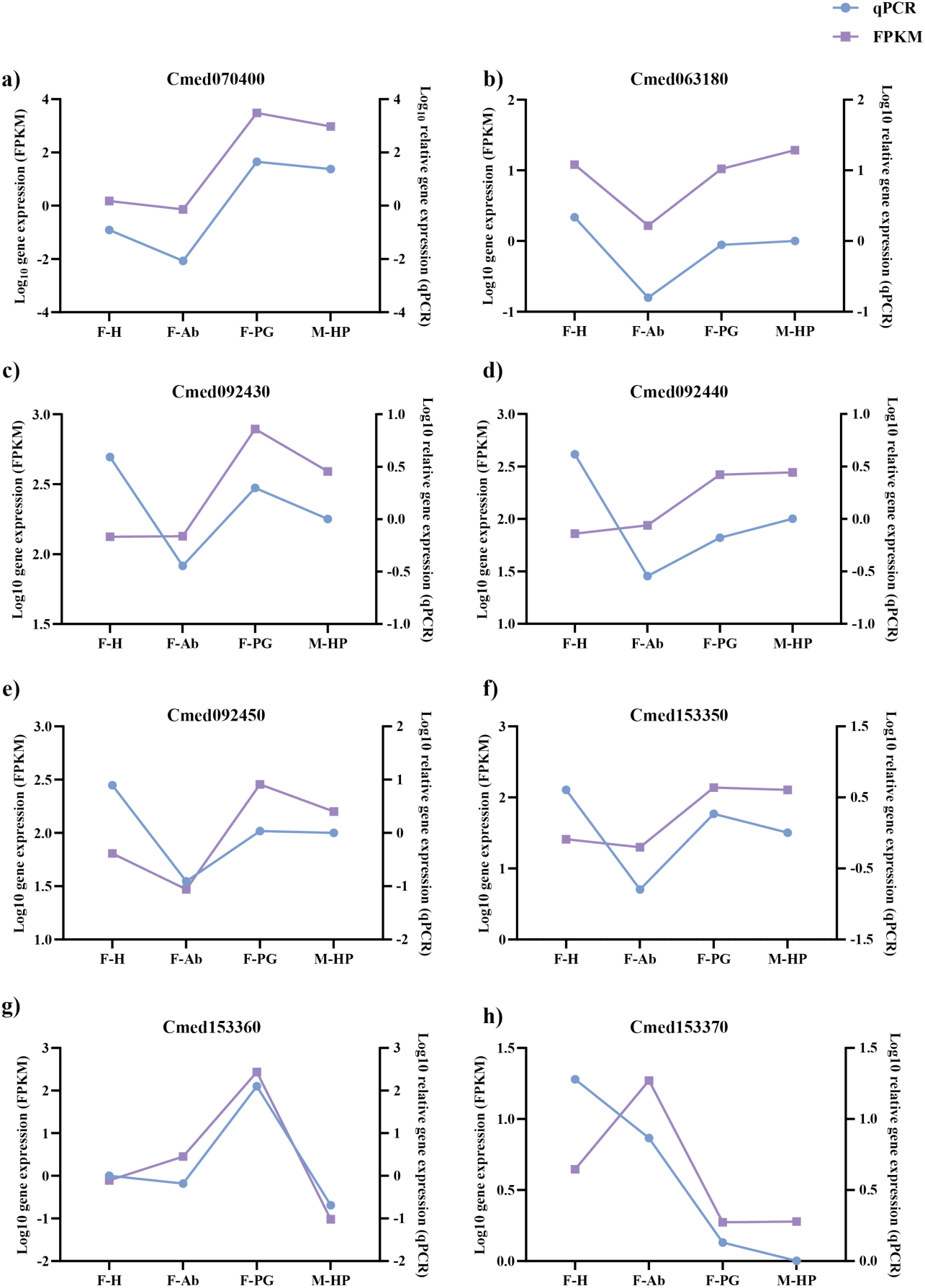
Validation of transcriptome-based expression profiles by quantitative PCR (qRT-PCR). Gene expression levels of selected fatty acyl desaturase (FAD) and elongase (ELO) genes in different tissues of *C. medinalis* were determined by qRT-PCR and compared with RNA-seq derived FPKM values. a) Cmed070400, b) Cmed063180, c) Cmed092430, d) Cmed092440, e) Cmed092450, f) Cmed153350, g) Cmed153360, h) Cmed153370.

### Sequence and phylogenetic analysis of FAD and ELO genes

Among the 12 FADs identified in the *C. medinalis* transcriptome, nine contained full-length open reading frames (ORFs). These include *Cmed009890, Cmed022650, Cmed054120, Cmed070400, Cmed077910, Cmed116530, Cmed116590, Cmed116600* and *Cmed138850*, encoding 333, 380, 359, 339, 301, 371, 376, 375, 303 aa-proteins with 999, 1140, 1077, 1017, 903, 1113, 1128, 1125, 909 nt ORFs, respectively. Sequence alignment revealed that all predicted FAD proteins possess the three conserved histidine boxes characteristic of membrane-bound desaturases (Fig. 5a). Phylogenetic analysis showed that *Cmed070400* clustered robustly within the Δ11 desaturase clade and shared the highest amino acid identity with *CsupYPAQ*, the major functional FAD involved in sex pheromone biosynthesis of Z11-16:acid in *Chilo suppressalis* (Fig. 6).

**Figure 5.**
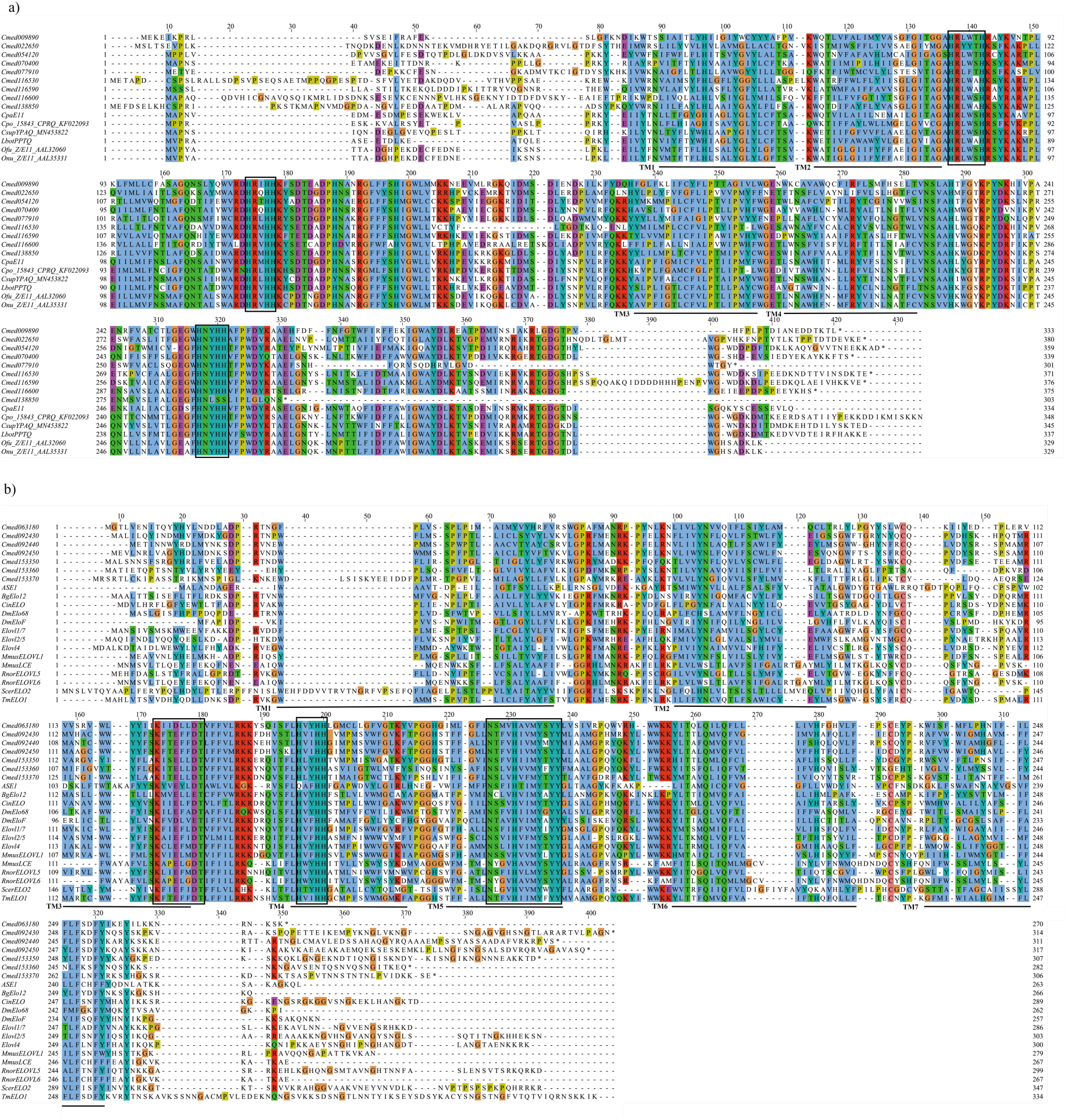
Amino acid sequence alignment of fatty acyl desaturases (FAD) and elongase (ELOs) candidates from *C. medinalis* and functionally characterized homologs. a) Multiple sequence alignment of candidate FAD proteins with representative FADs of known function. Predicted transmembrane helices are indicated by straight lines below the sequences. The three conserved histidine boxes are indicated by boxed regions. b) Multiple sequence alignment of candidate ELO proteins with representative elongases of known function. Predicted transmembrane helices are marked with straight lines. The conserved motifs characteristic of ELO proteins (KxxExxDT, HxxHH, and NxxVHxxMYxYY) are indicated by boxed regions.

**Figure 6.**
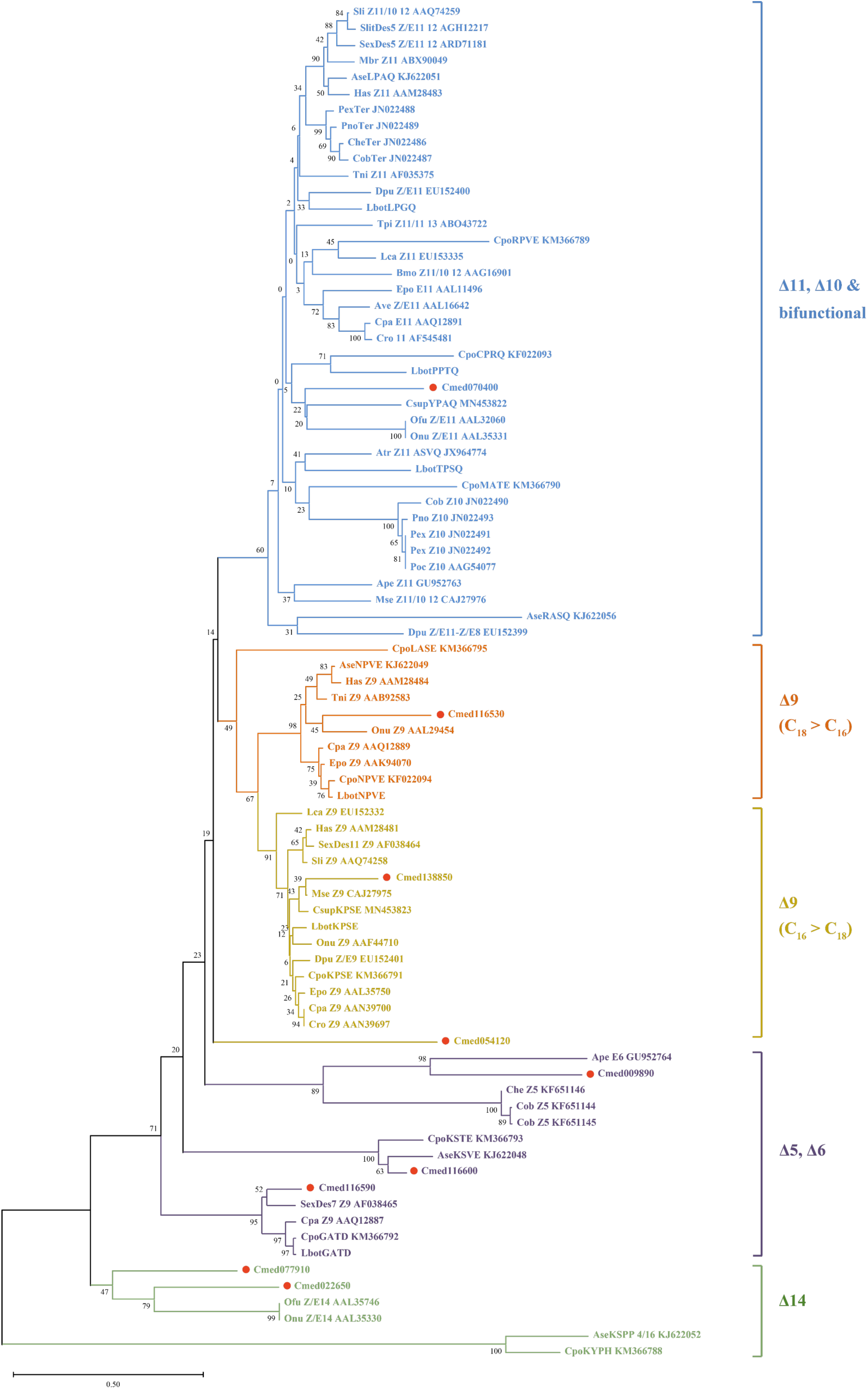
Phylogenetic analysis of fatty acyl desaturases (FADs). The tree was constructed using neighbor-joining (NJ) method with MEGA version 12.0.14. The predicted *C. medinalis* FAD candidates were indicated by red dots. Scale bar represents 0.50 amino acid substitutions per site. Branches are color-coded based on substrate specificity: blue, Δ11/Δ10/multifunctional desaturases; orange, Δ9 desaturases with a preference for C16 substrates; khaki, Δ9 desaturases with a preference for C18 substrates; purple, Δ5 and Δ6 desaturases; and green, Δ14 desaturases.

Seven elongase (ELO) candidates were identified with full-length ORFs: *Cmed063180*, *Cmed092430*, *Cmed092440*, *Cmed092450*, *Cmed153350*, *Cmed153360*, and *Cmed153370*. These genes encode proteins of 270, 314, 311, 317, 307, 259, and 306 aa-proteins amino acids, corresponding to ORFs of 810, 942, 933, 951, 921, 826, and 918 nt, respectively. Multiple sequence alignment revealed the presence of three conserved motifs (KxxExxDT, HxxHH, and NxxVHxxMYxYY), which are typical of ELO family proteins and are located within their central domains (Fig. 5b). The *C. medinalis* ELO-like proteins share moderate sequence identity (up to ∼40%) with characterized elongases such as mouse MmusELOVL1 (Table S1), supporting their classification as elongase-like enzymes. Phylogenetic analysis using functionally characterized ELO proteins from diverse taxa (including insects, vertebrates, yeast, and algae) showed that the *C. medinalis* ELO candidates cluster within the polyunsaturated fatty acid (PUFA) elongase subfamily (Fig. 7). This clustering suggests that these enzymes may participate in elongation of unsaturated fatty acyl substrates during pheromone precursor biosynthesis.

**Figure 7.**
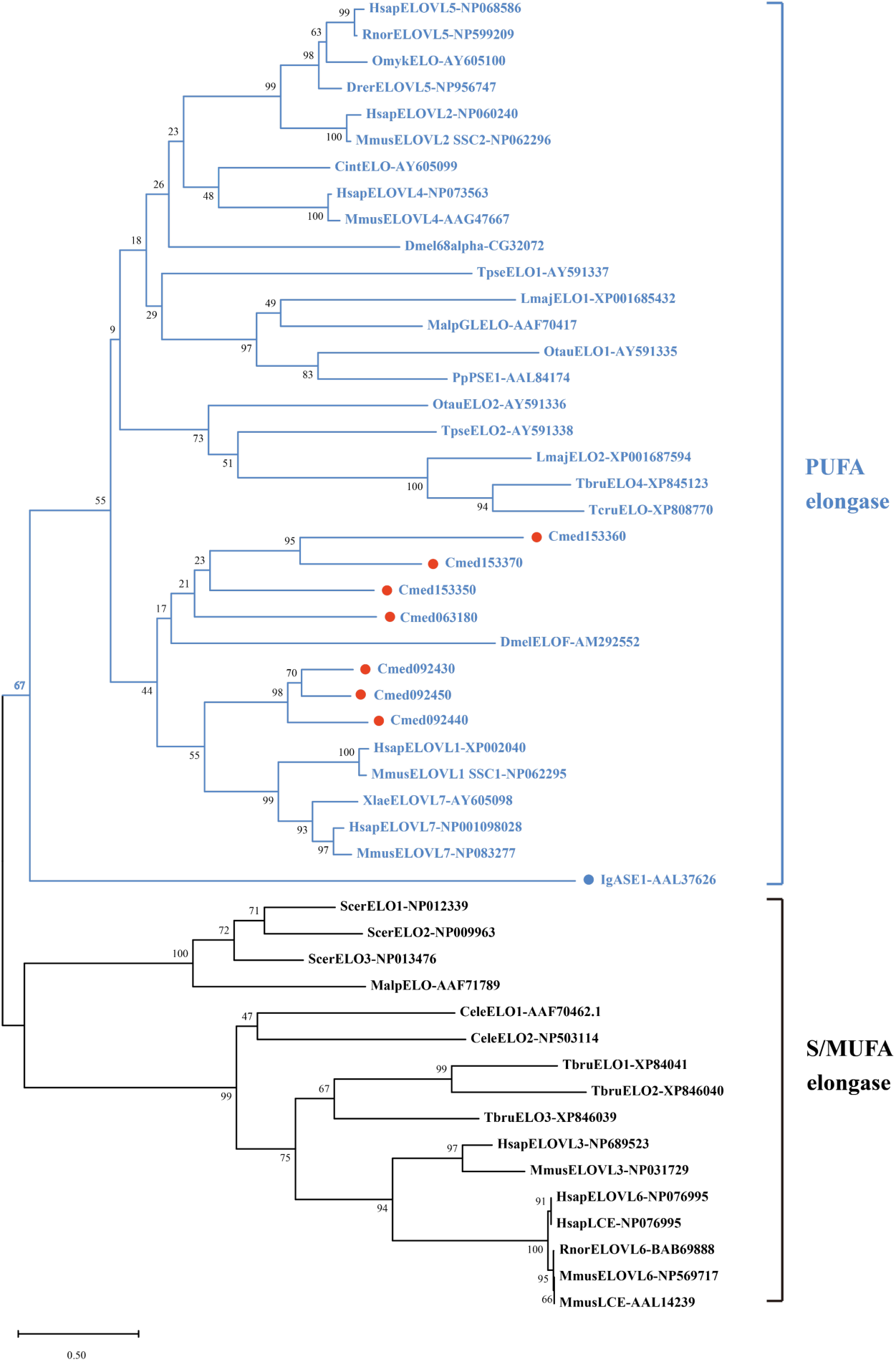
Phylogenetic analysis of elongases (ELO). The tree was constructed using neighbor-joining (NJ) method with MEGA version 12.0.14. Predicted ELO candidates from *C. medinalis* are indicated by red dots. The scale bar represents 0.50 amino acid substitutions per site. Branches are color-coded based on substrate specificity: red, polyunsaturated fatty acid (PUFA) elongases; black, monounsaturated and saturated fatty acid elongases.

### *Cmed070400* is involved in the desaturation during pheromone biosynthesis in *C. medinalis*

Heterologous expression of *C. medinalis* FAD candidates in *N. benthamiana* showed that only leaves expressing *Cmed070400* produced Z11-16:acid (Fig. 8c), whereas no such product was detected in leaves expressing other FAD candidates or in wild-type controls (Fig. 8). This result indicates that *Cmed070400* catalyzed the introduction of a cis double bond at Δ11 position of palmitic acid, generating Z11-16:acid, a key precursor in sex pheromone biosynthesis in *C. medinalis*. Consistent with this functional role, *Cmed070400* exhibited the highest expression level among desaturase genes in the pheromone gland (Fig. 3). qPCR further confirmed that its expression is significantly enriched in this tissue compared to other body parts (Fig. 4a). Together, these results support a central role of *Cmed070400* in pheromone precursor biosynthesis in *C. medinalis*.

**Figure 8.**
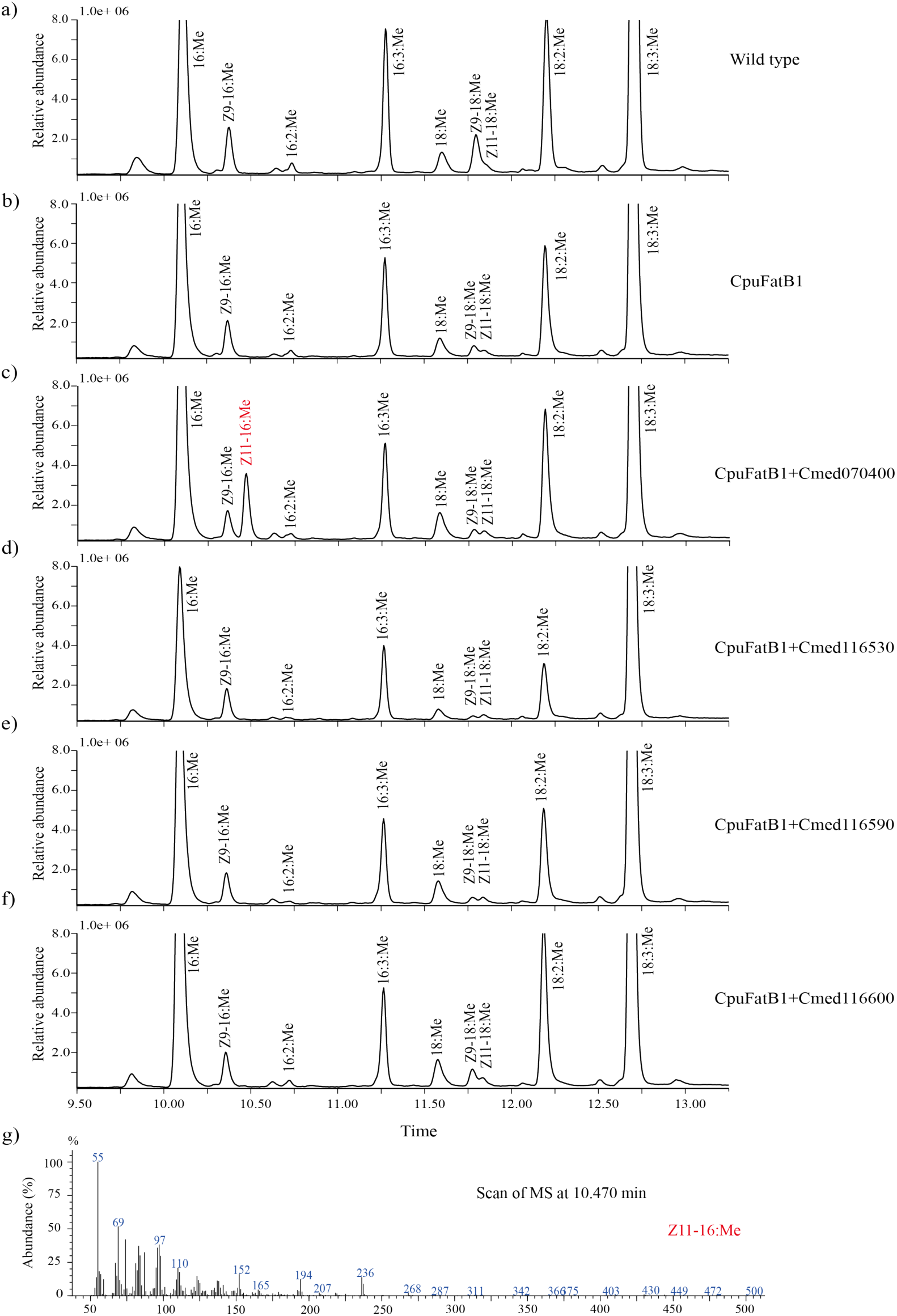
Functional characterization of FAD candidates by heterologous expression in *N. benthamiana*. Gas Chromatography-Mass Spectrometry (GC-MS) analysis of fatty acid methyl ester (FAME) profiles from a) wild type leaves, and leaves expressing b) *CpuFatB1*; c) *CpuFatB1*+*Cmed070400;* d) *CpuFatB1*+*Cmed116530;* e) *CpuFatB1+Cmed116590;* f) *CpuFatB1+Cmed116600*. Compounds derived from endogenous plant metabolism are shown in black, whereas products generated by the introduced desaturase are indicated in red. The internal standard (19:Me, methyl nonadecanoate) co-eluted with 16:2:Me (hexadecadienoic acid methyl ester) and is therefore not labeled separately. The compounds are shown in abbreviations: 16:Me, methyl palmitate; Z11-16:Me, (Z)-11-hexadecenoic acid methyl ester; Z9-16:Me, (Z)-9-hexadecenoic acid methyl ester; 16:3:Me, hexadecatrienoic acid methyl ester; 18:Me, methyl stearate; Z9-18:Me, (Z)-9-octadecenoic acid methyl ester; Z11-18:Me, (Z)-11-octadecenoic acid methyl ester; 18:2:Me, octadecadienoic acid methyl ester; 18:3:Me, octadecatrienoic acid methyl ester. i) Electron-ionization mass spectrum of the Z11-16:Me.

### Multiple ELOs exhibit fatty acyl chain elongation activity in a heterologous system, with *Cmed092440* showing the highest activity *Cmed092440*

Functional characterization of *C. medinalis* elongase (ELO) candidates in *N. benthamiana* demonstrated that several ELOs are capable of catalyzing fatty acyl chain elongation (Fig. 9–10). When expressed alone, the Δ11 desaturase *Cmed070400* produced a high level of Z11-16:acid, along with a minor amount of Z13-18:acid (Fig. 9b). Co-expression of Cmed070400 with individual ELO candidates resulted in differential production of Z13-18:acid. Notably, co-expression with *Cmed092440* led to a marked increase in Z13-18:acid accumulation compared with expression of *Cmed070400* alone (Fig. 9–10). Quantitative analysis showed that the conversion ratio of Z11-16:acid to Z13-18:acid increased more than fivefold in the presence of Cmed092440 (Fig. 10c), indicating enhanced elongation efficiency. Interestingly, *Cmed063180* exhibited a comparable conversion rate to *Cmed092440* (Fig. 10c), suggesting similar catalytic efficiency in terms of substrate conversion. However, the absolute production of Z13-18:acid was higher in plants expressing *Cmed092440*, indicating that *Cmed092440* achieves greater product accumulation under the tested conditions (Fig. 10b). Other ELO candidates displayed lower or negligible activity. Taken together, these results indicate that multiple elongases contribute to pheromone precursor elongation, with Cmed092440 showing a higher capacity for product formation in the heterologous system.

**Figure 9.**
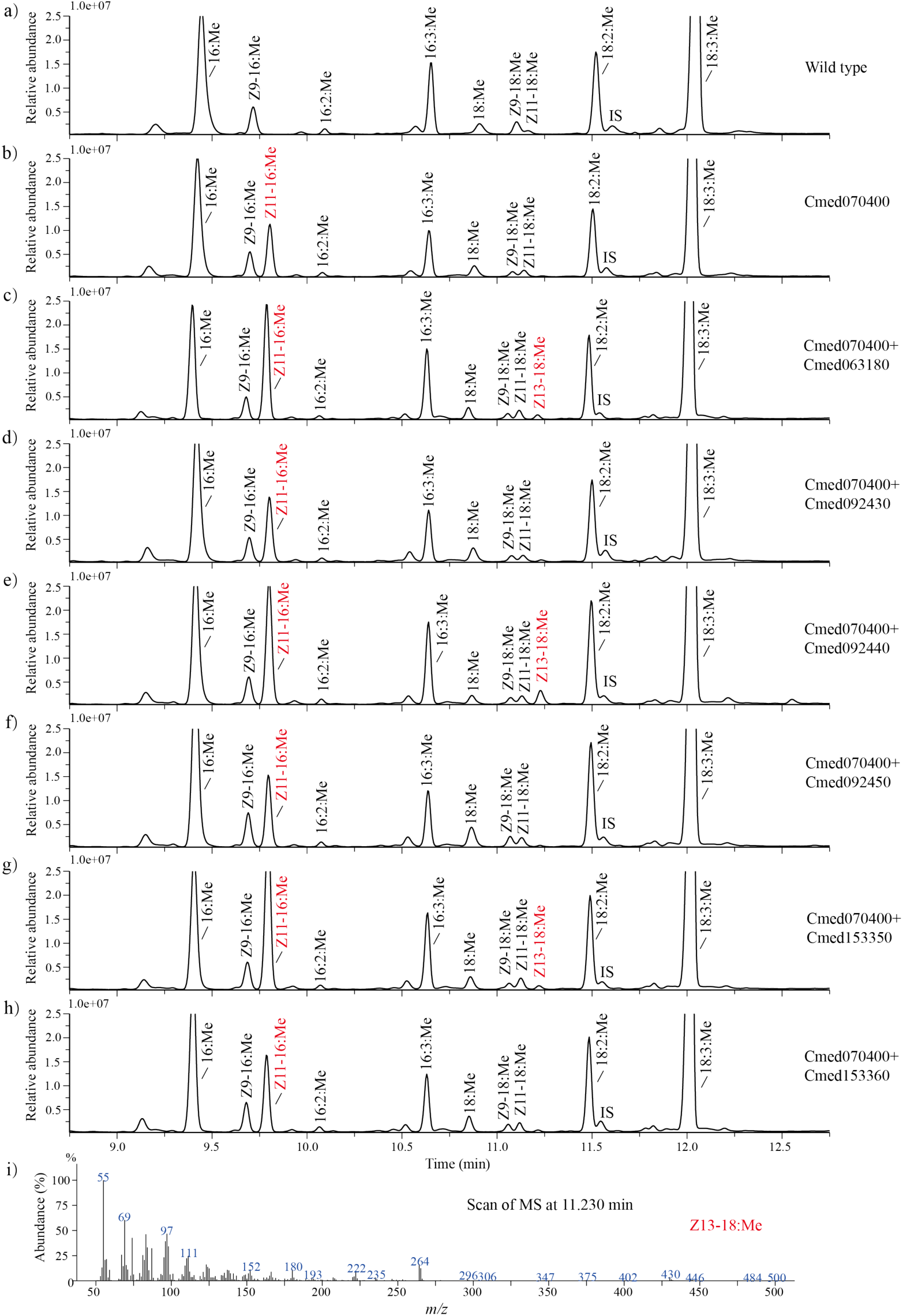
Functional characterization of ELOs candidates by heterologous expression in *N. benthamiana*. Gas Chromatography-Mass Spectrometry (GC-MS) analysis of fatty acid methyl ester profiles from a) wild type leaves, and the leaves expressing b) *Cmed070400*; c) *Cmed070400*+*Cmed063180*; d) *Cmed070400*+*Cmed092430*; e) *Cmed070400*+*Cmed092440*; f) *Cmed070400*+*Cmed092450*; g) *Cmed070400*+*Cmed153350*; h) *Cmed070400*+*Cmed153360*. Compounds derived from endogenous plant metabolism are shown in black, whereas products generated by the introduced desaturase are indicated in red. The compounds are shown in abbreviations: 16:Me, methyl palmitate; Z11-16:Me, (Z)-11-hexadecenoic acid methyl ester; Z9-16:Me, (Z)-9-hexadecenoic acid methyl ester; 16:2:Me (hexadecadienoic acid methyl ester) 16:3:Me, hexadecatrienoic acid methyl ester; 18:Me, methyl stearate; Z9-18:Me, (Z)-9-octadecenoic acid methyl ester; Z11-18:Me, (Z)-11-octadecenoic acid methyl ester; 18:2:Me, octadecadienoic acid methyl ester; 18:3:Me, octadecatrienoic acid methyl ester. The internal standard, 19:Me, is abbreviated as IS in the figure. i) Electron-ionization mass spectrum of the Z13-18:Me.

**Figure 10.**
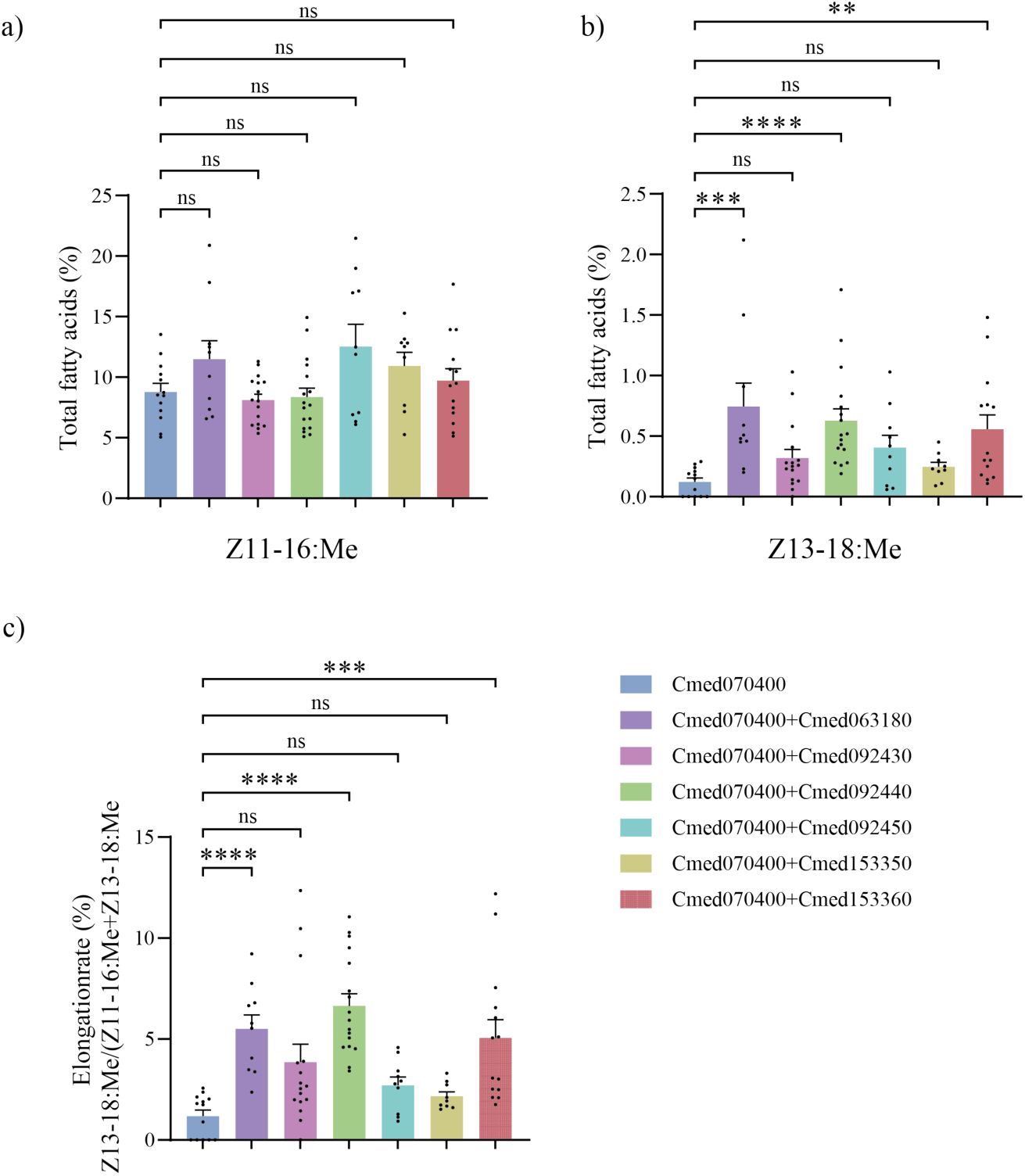
Statistical comparison of Z11-16:acid and Z13-18:acid production and elongation efficiency among different gene combinations in *N. benthamiana*. a) Production of Z11-16:acid, b) Production of Z13-18:acid, c) Conversion rate of Z11-16:acid to Z13-18:acid in leaves expressing *Cmed070400* alone or co-expressing *Cmed070400* with different ELO candidates. All measurements were performed on fatty acid methyl esters (FAMEs). Data are presented as mean ± standard error of the mean (SEM), with each independent experiment including 3–5 biological replicates (total n ≥ 10). For a), data are presented as the mean ± S.E.M. and analyzed by one-way ANOVA. For b) and c), which violated normality assumptions, the Kruskal-Wallis test followed by Dunn’s post-hoc test was used. Data normality reports are provided in Supplementary Table S3–5. *P* < 0.05 was accepted as statistically significant. The error bars represent the standard error of the mean (S.E.M.).

## Discussion

Sex pheromone biosynthesis in Lepidoptera relies on a coordinated sequence of enzymatic reactions that determine both the chemical structure and species specificity of pheromone signals. In this study, our findings define a desaturation–elongation sequence underlying pheromone precursor formation in *C. medinalis*, in which Δ11 desaturation generates the key intermediate Z11-16:acid that is subsequently extended by elongases to yield Z13-18:acid (Fig. 1D), the direct precursor of the major pheromone component (Fig. S3). This intermediate is further reduced by FAR to the corresponding alcohol (Z13-18:OH); in some moths species, this alcohol itself functions as the pheromone (e.g., *Cydia pomonella*, *Lampronia capitella*), whereas in more common cases it undergoes a final oxidation to the aldehyde (Z13-18:Ald), which constitutes the major pheromone component in *C. medinalis* [26, 44–46]. Despite extensive biochemical evidence supporting this final oxidation step for aldehyde formation [47, 48], the responsible oxidase remains unidentified at the gene level. Our work therefore resolves the upstream biosynthetic framework while highlighting a persistent gap at the terminal step, although four candidate ADHs identified in this study may warrant further functional investigation (Table 2).

Among the functionally characterized genes in the present study, multiple ELOs were found to catalyze Z11-16:acid to form Z13-18:acid, implying that pheromone precursor elongation in *C. medinalis* may be mediated by a set of enzymes rather than a single dedicated elongase. In particular, *Cmed092440*, *Cmed063180*, and *Cmed153360* all exhibited elongation activity in the heterologous expression system, although their catalytic outputs differed (Fig. 10). Expression of *Cmed070400* alone in *N. benthamiana* resulted in the accumulation of Z11-16:acid and trace amounts of Z13-18:acid, likely reflecting low background activity of endogenous plant elongases (Fig. 9). In contrast, co-expression of *Cmed070400* with individual

elongases led to increased production of Z13-18:acid, with *Cmed092440* consistently yielding the highest product accumulation. Notably, *Cmed063180* displayed a comparable conversion efficiency but lower overall product levels, suggesting that elongases may differ in catalytic efficiency, expression level, or stability in the heterologous system. The deduced proteins of the identified *C. medinalis* elongases contain the conserved HxxHH histidine-rich motif, which is proposed to function as an iron-chelating site involved in electron transfer during oxygen-dependent redox reactions [49]. In addition, these proteins possess seven predicted transmembrane domains, a structural feature conserved among elongases characterized in yeast and mammals [28, 29, 50] (Fig. 5b). The presence of these hallmark catalytic motifs and membrane-associated domains, together with their comparable expression levels in the pheromone gland, strongly supports the annotation of these genes as bona fide elongases with roles in fatty acyl chain elongation during pheromone biosynthesis. Although this approach provides robust evidence for catalytic capability, the *in vivo* functions of these elongases in *C. medinalis* are likely influenced by additional factors, including substrate availability, subcellular localization, and interactions with other components of the native biosynthetic machinery. Therefore, further in vivo functional validation will be required to fully elucidate the physiological roles and relative contributions of individual elongases in pheromone biosynthesis.

Functional characterization of insect elongases remains limited, in part due to their relatively low sequence similarity to previously annotated homologs. For example, Cmed092440 shows highest identity to mouse ELOVL1, but only at 43.62% (Table S1), highlighting the evolutionary divergence of this enzyme family. Consistent with observations across taxa, elongases often exhibit broad substrate specificity. For instance, *MmusELOVL1* primarily elongates PUFAs but can complement yeast mutants lacking elongase *elo3* activity toward SFA/MUFAs [50]. While *RnorELOVL5* knockdown Rat line showed decreased elongation of MUFA of 16:1,n-7 [51]. Similarly, *Drosophila Elo68α* and *EloF* display broad substrate ranges. The former one can carry out elongation of Z9-14:acid, Z9-16:acid and Z11-16:acid [33] and the later one can elongate very long chains of saturated and unsaturated fatty acids from C19 to C30 [34]. In addition, Zebrafish *Danio rerio* elongase *DrerELOVL5* not only lengthens the range of C18, C20 and C22 PUFAs but also able to elongate S/MUFAs [52]. In line with this functional flexibility, the phylogenetic placement of *C. medinalis* elongases within the PUFA clade does not preclude activity toward monounsaturated substrates such as Z11-16:acid. The absence of additional elongated products in the plant system likely reflects limited availability of alternative substrates rather than strict enzymatic specificity.

Plant transient expression represents a robust and discovery-oriented platform for functional characterization of elongases. For example, the algal elongase IgalASE1 elongated 18:2 and 18:3 to 20:2 and 20:3 in *N. benthamiana* (Figure S1), consistent with its activity in *S. cerevisiae* [53]. Although transient expression of insect genes in plant systems has been successfully applied for pheromone production [2], its use in dissecting pheromone biosynthetic enzyme functions relatively limited. Compared with yeast-based systems, plant assays avoid interference from exogenously supplied fatty acids and provide a lipid environment that more closely reflects native substrate pools. This distinct metabolic context can reveal minor or context-dependent enzymatic activities that may not be detectable in yeast. For instance, *CsupFAR2* produced additional low-abundance products from α-linolenic acid in *N. benthamiana* compared with yeast [54]. Here, the plant system enabled a clear reconstruction of the desaturation-then-elongation logic leading to Z13-18:acid and revealed a low endogenous elongation background, thereby providing a useful baseline for evaluating the activity of *C. medinalis* elongases. Together, these findings highlight the utility of *N. benthamiana* as a complementary platform for elucidating the functional diversity of pheromone biosynthetic genes.

Functional characterization of *C. medinalis* elongases also provides broader insights into the evolution of pheromone biosynthesis in Lepidoptera. Current databases document pheromones and attractants in Lepidoptera (https://lepipheromone.sakura.ne.jp/lepi_phero_list_eng.html), showing more than 300 Lepidoptera species, ranging from Tineoidea to Noctuoidea that use pheromone or attractant components with the fatty acyl chain of Z11-16 [54, 55] (Fig. 11). The diversification of pheromone signals is thought to play a key role in reproductive isolation and speciation, with enzymatic innovations, particularly among desaturases, driving structural variation in pheromone components [5, 55, 56]. Our findings suggest that elongases have likely evolved in parallel with desaturases to shape pheromone diversity. The phylogenetic analysis suggests that lepidopteran ELOs were coopted for pheromone biosynthesis at least 150 million years ago before the divergence of the superfamily Tineoidea and the more advanced Lepidoptera. Most of the Tineoidea spp. for which pheromones were identified use unusual Type I pheromone compounds, Δ3,Z13-18 or Δ2,Z13-18 alcohols, acetates or aldehydes, which are hypothesized to be biosynthesized by Δ11 desaturation of palmitic acid, followed by the formation of the second double bond in connection with the chain elongation [5, 6]. Similarly in Sesioidea, 108 identified pheromones/attractants may involve an elongase to produce the carbon skeletons Δ2/3,Z13-18 of their major pheromone components from Z11-16:acid (Fig. 11). Thus, our elongase finding provides the missing genetic lever for a lineage-restricted chemical trajectory, linking enzyme identity to macroevolutionary signal patterns.

**Figure 11.**
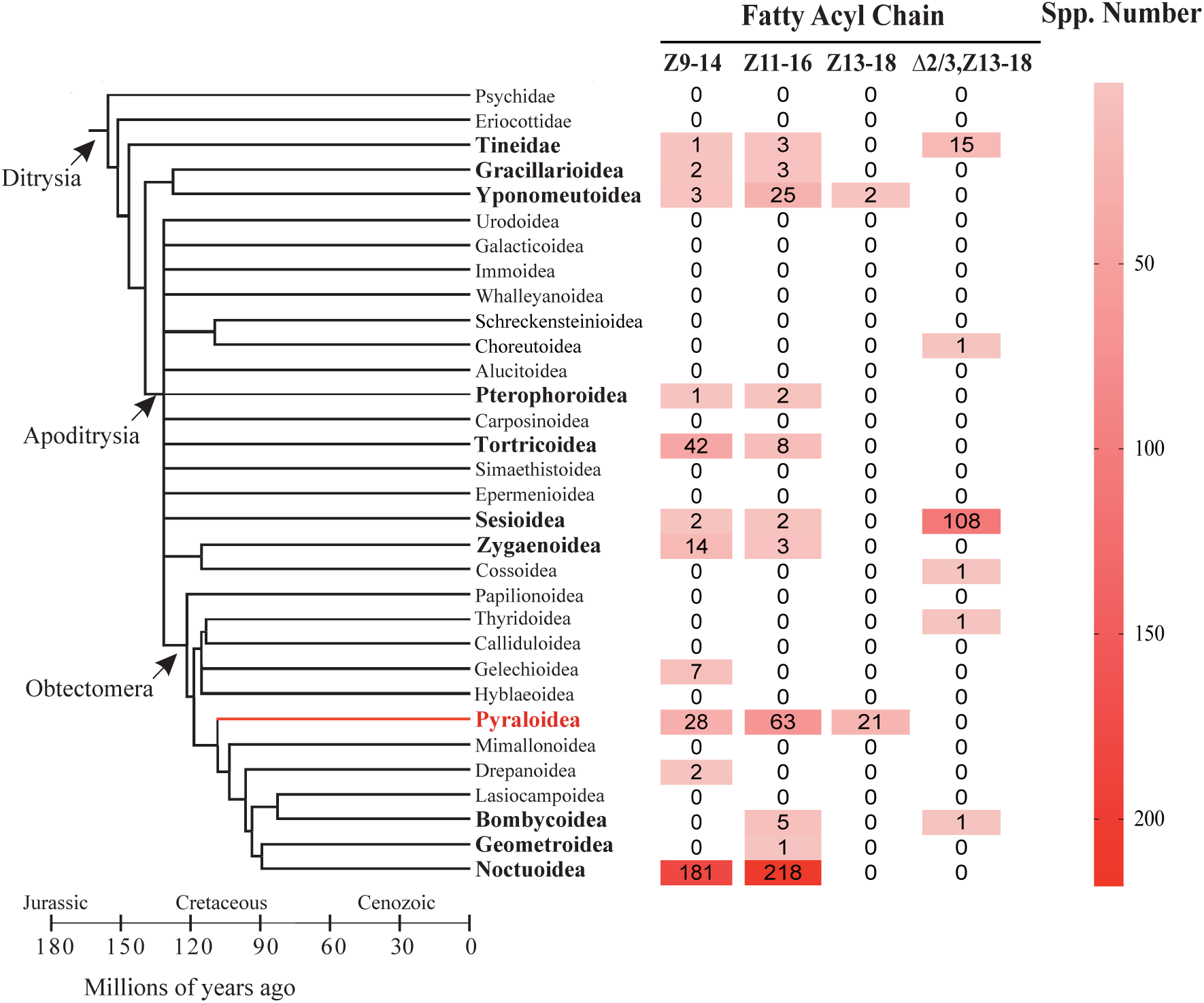
Fatty acyl chain of pheromone and attractant components mapped on to ditrysian Lepidoptera superfamilies. Numbers after taxa indicate reported number of species using corresponding fatty acyl chains in their pheromones and attractants in each superfamily. Z9-14 refers to the fatty acyl chain with a chain length of 14-carbon atoms and a double bond at Δ9 position in ‘Z’ configuration. Z11-16 refers to the fatty acyl chain with a chain length of 16-carbon atoms and a double bond at Δ11 position in ‘Z’ configuration. Z13-18 refers to the fatty acyl chain with a chain length of 18-carbon atoms and a double bond at Δ13 position in ‘Z’ configuration. Δ2/3,Z13-18 refers to the fatty acyl chain with a chain length of 18-carbon atoms and one double bond at Δ2 or Δ3 position in ‘Z’ or ‘E’ configuration, and the other double bond at Δ13 position in ‘Z’ configuration. The number of the fatty acyl chain is counting from the internet database on https://lepipheromone.sakura.ne.jp/lepi_phero_list_eng.html [72]

Notably, elongation of Z11-16 to Z13-18 appears to be a lineage-restricted biochemical trajectory, predominantly observed in Yponomeutoidea and Pyraloidea (Fig. 11). In several pyraloid species, including the sugarcane striped borer *Chilo auricilius* [57], the Mexican rice borer *C. loftini* [58], and *C. medinalis* [59, 60], Z13-18-derived compounds constitute their major pheromone components. In this context, the identification of Cmed092440 and other active elongases provides a molecular explanation for this +2 carbon extension pathway and links enzyme function to lineage-specific pheromone evolution. More broadly, our results suggest that diversification within the elongase gene family, together with variation in catalytic efficiency and substrate utilization, may represent an additional layer of regulation contributing to species-specific pheromone signals.

Beyond evolutionary implications, these findings also have practical significance for biotechnological applications. While we successfully demonstrate the production of Z13-18:acid through heterologous expression of *C. medinalis* elongases, further optimization will be required to enhance production yields for industrial applications. Potential strategies include codon optimization of these ELO genes and co-expression of multiple active ELOs, such as *Cmed092440* with *Cmed153360* and *Cmed063180* which may act synergistically to improve flux through the elongation pathway. A particularly promising approach for improvement involves reconstruction the native elongation machinery. In eukaryotes, fatty acid elongation is not mediated by a single enzyme but by a multi-component complex located to the endoplasmic reticulum, comprising a β-ketoacyl-CoA synthase (KCS), a β-ketoacyl-CoA reductase (KCR), a β-hydroxyacyl-CoA dehydratase (HCD), and an enoyl-CoA reductase (ECR) [61, 62, 63]. The spatial organization of these enzymes facilitates substrate channeling, thereby enhancing catalytic efficiency and minimizing loss of intermediates to competing pathways [57, 61, 64–67]. Reconstituting such a system in heterologous hosts, potentially through co-expression and synthetic scaffolding of elongase-associated enzymes, may enable the formation of a functional metabolon. We hypothesize that this approach could substantially improve catalytic efficiency and stability of the elongation process, ultimately increasing the yield of Z13-18:acid.

## Conclusions

In summary, by resolving the two upstream, rate-setting steps—Δ11 desaturation and +2C elongation—this study assigns a causal enzymatic basis to the major pheromone precursor in *C. medinalis*. Our results identify elongases as key determinants of pheromone signal architecture alongside desaturases and establish a functional desaturation–elongation framework underlying Z13-18 biosynthesis. The identification of multiple active elongases, with *Cmed092440* showing the highest productive capacity, provides a minimal and transferable gene set for pathway reconstruction in plant systems. This work also validates *N. benthamiana* as a robust platform for functional interrogation of pheromone biosynthetic enzymes. The terminal oxidation step remains unresolved at the genetic level, representing a critical target for future study. Linking variation in Cmed092440 and Cmed070400 to in vivo pheromone production and population-level signal divergence will be essential to fully understand pathway regulation. Overall, this study advances both the mechanistic understanding of moth pheromone biosynthesis and the development of plant-based platforms for sustainable production of Z13-18-derived compounds.

## Materials and Methods

### Insect rearing and tissue collection

The *C. medinalis* specimens used were originally collected in China (22.80°N, 113.95°E) and have been successively reared on an artificial diet at 26 °C under a long-day cycle (16L, 8D). All experiments were carried out under the above conditions. Pupae were sexed and separately held in plastic-screen cages. After emergence, female and male adults were kept separately under the same conditions as above and provided with 10% honey solution as food. Moth age was defined as day 0 on the day of emergence, and as day 1, day 2, and so on for subsequent days.

Fourteen tissue types were collected from adult *C. medinalis* at 2-3 days post-emergence during the scotophase (2-5 hours after onset). To ensure biological reproducibility and minimize individual variation, each sample comprised a pooled mixture of approximately 30 individuals to overcome the constraints of low biomass. Specifically, the samples included heads, thoraxes, abdomens, legs, wings, and antennae from both sexes, as well as female pheromone glands (PG) and male hair-pencils (HP). All samples were flash-frozen in liquid nitrogen and subsequently used for RNA extraction.

### Total RNA extraction and transcriptome sequencing

Total RNA was extracted from 14 tissue types dissected from more than 20 adult moths using the TRIzol reagent, following the manufacturer’s instructions. RNA concentration and purity were determined using a NanoDrop spectrophotometer, with samples showing A260/280 ratios between 1.8-2.0 considered acceptable. Qualified RNA samples were sent to Novogene Bioinformatics Technology Co., Ltd. (Beijing, China) for reference-based transcriptome sequencing, on an Illumina NovaSeq X Plus platform with a paired-end 150 bp (PE150) configuration. To ensure comprehensive transcriptome characterization, both reference genome-guided and *de novo* assembly strategies were employed. For *de novo* assembly, clean reads were assembled using Trinity. For the reference-guided assembly, clean reads were mapped to the reference genome *C. medinalis* (GenBank: GCA_014851415.1) using HISAT2. Tissue dissection and RNA sequencing were performed independently twice to generate biological replicates.

### Validation of transcriptome-derived tissue expression profiles by qRT-PCR

Quantitative real-time PCR (qRT-PCR) was performed to validate the expression levels of selected genes. First-strand cDNA was synthesized from 500 ng of total RNA using a reverse transcription kit (HiScript Ⅲ RT SuperMix for qPCR (+gDNA wiper), Vazyme, China) according to the manufacturer’s instructions. qRT-PCR reactions were carried out in a 20 μL reaction mixture containing 10 μL of AceQ Universal SYBR qPCR Master Mix (Vazyme, China), 0.5 μL of each primer (10 μM), 9 μL of cDNA template. The amplification condition was as following: initial denaturation at 95 °C for 5 min, followed by 40 cycles of 95 °C for 10 s and 60 °C for 30 s, with a final melting curve analysis to confirm specificity. Due to the extreme difficulty of sample collection and the minute size of the target tissues, each analyzed sample represented a single biological replicate consisting of a pooled mixture of approximately 30 individuals to minimize individual variation. To ensure the highest possible reliability in the absence of traditional biological replicates, all qRT-PCR reactions were strictly run in triplicate (technical replicates) with rigorous quality control, including the verification of single peaks in melting curves. Each sample was analyzed in triplicate. Relative gene expression levels were calculated using the 2^(-ΔΔCt) method, with PGK as the internal reference gene.

### Sequences and phylogenetic analysis

The open reading frames (ORFs) of ELO candidates were obtained from the *C. medinalis* reference genome (GenBank: GCA_014851415.1). These sequences, along with functionally characterized homologs from NCBI GenBank, were aligned using the MUSCLE algorithm implemented in MEGA v12.0.14[68]. A phylogeny was then constructed using the Neighbor-Joining method. The best-fit nucleotide substitution model was automatically selected by the software, and branch support was assessed with 1000 bootstrap replicates. For the analysis of pheromone fatty acyl chain in different lepidopteran superfamilies, the phylogenetic tree of Lepidoptera was adapted and modified from Löfstedt et al. 2016 [5].

### Full-Length cDNA Cloning Using Rapid Amplification of cDNA Ends (RACE)

Initial attempts to amplify the *Cmed153360* gene via standard PCR using cDNA as the template were unsuccessful. Given its unusually long sequence compared to other elongase candidates, we suspected that the reference genomic sequence might be artifactual. Therefore, rapid amplification of cDNA ends (RACE) was employed to obtain the authentic full-length cDNA. RACE was performed using the HiScript-TS 5’/3’ RACE Kit (Vazyme, Nanjing, China), and the resulting amplicons were cloned (pCE3 Blunt vector, Vazyme) and Sanger-sequenced. The internal, 5’, and 3’ RACE fragments were assembled to reconstruct the complete coding sequence. The final sequence exhibited high homology with other elongase candidates. Furthermore, PCR primers designed against this assembled sequence successfully amplified *Cmed153360*, yielding a product identical to our RACE assembly, thereby confirming its accuracy.

### Cloning of gene candidates for functional assay

PCR amplification of each gene candidate was performed using cDNA as template with a pair of specific primers (Table S2) with attB1 and attB2 sites incorporated on an ETC811 thermal cycler (ETC, Hangzhou, China) using KOD One TM PCR Master Mix-Blue- (Takara, Japan). Cycling parameters were as follows: an initial denaturing step at 94 °C for 3min, 38 cycles at 98 °C for 5 s, 55 °C for 10 s, 68 °C for 30s, followed by a final extension step at 68 °C for 10 min. For several candidate genes: Cmed009890, Cmed022650, Cmed054120, Cmed077910, Cmed138850, Cmed153370, no PCR product of the expected size was observed, despite the use of optimized cycling conditions and multiple primer sets. The PCR products were subjected to agarose gel electrophoresis and purified using the GeneJET Gel Extraction Kit (Thermo Scientific™). Then the ORFs were cloned into the pDONR221 vector in present of BP clonase (Life Technologies) to generate entry clone. After the entry clone for each ORF was confirmed by sequencing with M13F and M13R primers to be correct, it was cloned into plant expression vector pXZP393 by LR reaction (Invitrogen). The resulting expression clones were analyzed by sequencing.

### *N. benthamiana* material and growth condition

The wild type *N. benthamiana* plants for the *Agrobacterium* infiltration were grown in the greenhouse under 16 h/8 h light conditions. Growth temperature and relatively humidity in greenhouse were set at 24 °C/18 °C in day/night and 40%, respectively.

### Agrobacterium infiltration of N. benthamiana

The viral silencing suppressor protein gene *P19* (WAK97598.1) was introduced into GV3101 strain to inhibit transgene silencing[69]. Additionally, the plastid acyl-ACP thioesterase gene *CpuFatB1* (AGG79283.1) from *Cuphea pulcherrima* was transformed into the same strain. This gene encodes an enzyme capable of hydrolyzing acyl-chain-ACP to release free saturated C16 fatty acids [3, 70], thereby providing a substrate for the downstream desaturase and elongase. The elongase gene *IgalASE1* was also transformed as a positive control. The *Agrobacterium* strains harboring individual gene constructs were cultured overnight in LB medium with appropriate antibiotics at 30 °C. The cultures were then induced with 100 μM acetosyringone for 2-3 h, pelleted, and resuspended in infiltration buffer (5 mM MgCl2, 5 mM MES, 100 µM acetosyringone, pH 5.7). The cultures were then induced with 100 μM acetosyringone for 2–3 h, pelleted, and resuspended in infiltration buffer (5 mM MgCl₂, 5 mM MES, 100 µM acetosyringone, pH 5.7). Then each *Agrobacterium* culture was adjusted to an OD600 of 0.2 in a total volume of 20 mL. The suspensions were infiltrated into the abaxial surface of five fully expanded leaves from five-week-old *N. benthamiana* plants. Infiltrated plants were maintained in a growth chamber for four days until analysis.

### Lipid extraction and preparation

Approximately 300 mg of *N. benthamiana* leaf tissue was randomly collected and transferred into a 4 mL glass tube. The sample was supplemented with 1 mL of 2% sulfuric acid in methanol and 3.12 μg of the internal standard methyl nonadecanoate (19:Me), followed by incubation at 90 °C for 1 h to facilitate methanolysis. After incubation, 1 mL of water and 1 mL of heptane were added, and the mixture was vigorously vortexed. The samples were then centrifuged at 2,000 rpm for 2 min to achieve phase separation. Finally, approximately 1 mL of the upper heptane phase, containing fatty acids as their corresponding methyl esters, was collected and transferred to a clean glass vial for GC–MS analysis.

### Gas chromatography/mass spectrometry (GC/MS)

Fatty acid methyl esters (FAMEs) were analyzed using a Shimadzu GC-MS-TQ8050NX triple quadrupole mass spectrometer equipped with a polar BPX-70 column (25 m × 0.22 mm i.d. × 0.25 μm film thickness; Trajan Scientific and Medical, Victoria, Australia). Helium was used as the carrier gas at an average linear velocity of 46.2 cm s⁻¹. The injector was operated in splitless mode at 250 °C. The oven temperature program was as follows: initial temperature of 80 °C for 1 min, increased to 230 °C at a rate of 10 °C min⁻¹, and held at 230 °C for 10 min. Mass spectra were acquired in full-scan mode over an m/z range of 50–500. Compounds were identified by comparison of retention times and mass spectra with those of authentic standards.

### Statistical analysis

Data are presented as mean ± standard error of the mean (SEM), based on three independent experiments, each comprising 3–5 biological replicates (total n ≥ 10). Statistical analyses were conducted using GraphPad Prism version 10.6.0 [71]. For datasets meeting normality assumptions, one-way analysis of variance (ANOVA) was applied. For datasets that did not meet normality assumptions, the nonparametric Kruskal–Wallis test followed by Dunn’s post hoc test was used. A significance threshold of P < 0.05 was applied. Data normality assessments are provided in Supplementary material.

## Abbreviations

C16: palmitic acid
C18: stearic acid
C22: docosanoic acid
C24: tetracosanoic acid
20:0: eicosanoic acid
18:2: octadecadienoic acid
18:3: octadecatrienoic acid
20:2: eicosadienoic acid
20:3: eicosatrienoic acid
Z11-16:acid: (Z)-11-hexadecenoic acid
Z13-18:acid: (Z)-13-octadecenoic acid
Z13-18:OH: (Z)-11-octadecen-1-ol
19:Me: methyl nonadecanoate
PG: pheromone glands
HP: hair-pencils
ORF: open reading frames
ELO: fatty acyl elongases
FAS: fatty acid synthase
ACC: acetyl-CoA carboxylase
FAD: fatty acyl desaturases
ACO: acyl-CoA oxidases
FAR: fatty acyl reductase
ADH: alcohol dehydrogenases
KCS: β-ketoacyl-CoA synthase
KCR: β-ketoacyl-CoA reductase
HCD: β-hydroxyacyl-CoA dehydratase
ECR: enoyl-CoA reductase
PUFA: polyunsaturated fatty acid
SFA: saturate fatty acid
MUFA: monounsaturated fatty acid

## Declarations

### Ethics approval and consent to participate

Not applicable.

### Consent for publication

Not applicable.

### Availability of data and materials

All data generated or analysed during this study are included in this manuscript and its supplementary information files. Transcriptome data have been deposited in China National National Center for Bioinformation (BioProject number: PRJCA050327, accession number: F_H:CRA034032,F_T:CRA034033,F_Ab:CRA034034,F_L:CRA034035,F_W:CRA034036, F_A:CRA034037,F_PG:CRA034038,M_H:CRA034491,M_T:CRA034040,M_Ab:CRA0340 41,M_L:CRA034042,M_W:CRA034043,M_A:CRA034044,M_HP:CRA034045).

### Authors’ contributions

YHX and LLZ conceived the study. XYL, KXW and LYC carried out insect collection and tissue dissection. YHX and LYC performed vector design and sequencing. LYC and XYL performed transcriptome analysis. LYC performed gene functional assays. LYC performed plant cultivation and sample analysis. YHX, LLZ, SLD, HTT and XF provided technical guidance and suggestions on experimental strategies. YHX and LYC drafted the manuscript. All authors contributed to editing the draft. The authors read and approved the final manuscript.

## Acknowledgements

We thank Peng He and Ya-Nan Zhang for valuable discussions. We thank Meng-Ru Liu for excellent technical support. We also thank Pherobio Technology Co., Ltd. (China) for collaboration and for providing materials and infrastructure that supported the execution of this study.

