## Supplementary materials for "Integrative Transcriptomic and Functional Analysis Reveals Fatty Acyl Elongases Involved in Sex Pheromone Biosynthesis in Rice Leaffolder, *Cnaphalocrocis medinalis* (Lepidoptera: Pyraloidea)"

**Table S1. Identity of *C. medinalis* ELO-like proteins to Drosophila 68α, Drosophila EloF, Ciona Elo, Isochrysis ASE1, mouse ELOVL1 (SSC1), mouse ELOVL6 (LCE), rat ELOVL5, rat ELOVL6 and yeast S.c. ELO2 elongases.**

|  | DmElo68α | DmEloF | CinELO | IgASE1 | MmusELOVL1 | MmusLCE | RnorELOVL5 | RnorELOVL6 | ScerELO2 |
| --- | --- | --- | --- | --- | --- | --- | --- | --- | --- |
| Cmed063180 | 30.71% | 30.86% | 38.24% | 19.78% | 36.43% | 20.73% | 31.37% | 20.73% | 23.08% |
| Cmed092430 | 34.08% | 30.12% | 36.64% | 25.46% | 41.49% | 21.17% | 35.08% | 21.74% | 19.81% |
| Cmed092440 | 34.83% | 31.66% | 35.74% | 24.35% | 43.62% | 22.99% | 33.99% | 22.99% | 20.31% |
| Cmed092450 | 34.21% | 31.66% | 37.11% | 23.90% | 40.43% | 23.555 | 34.43% | 23.55% | 21.98% |
| Cmed153350 | 34.08% | 31.01% | 36.99% | 22.88% | 38.79% | 23.19% | 30.36% | 23.19% | 18.52% |
| Cmed153360 | 26.22% | 29.96% | 31.25% | 17.71% | 29.89% | 19.27% | 25.87% | 19.27% | 20.53% |
| Cmed153370 | 25.47% | 31.01% | 32.42% | 19.56% | 29.79% | 21.74% | 27.99% | 22.10% | 19.02% |

**Table S2. Primers used in this study.**

| Gene abbreviation | Primer sequence (5’-3’) |
| --- | --- |
| attB1_Cmed92430 For | GGGGACAAGTTTGTACAAAAAAGCAGGCTTAATGGCGCTAATACTTCAATATATAAAC |
| attB2_Cmed092430 Rev | GGGGACCACTTTGTACAAGAAAGCTGGGTTTCAGTTGCCGGCCGGCAGCACGG |
| attB1_Cmed092440 For | GGGGACAAGTTTGTACAAAAAAGCAGGCTTAATGGAGACCATAAACAACTGGTAC |
| attB2_Cmed092440 Rev | GGGGACCACTTTGTACAAGAAAGCTGGGTTCTAGGACACGGGCCGCTTGCGCACG |
| attB1-Cmed092450 For | GGGGACAAGTTTGTACAAAAAAGCAGGCTTAATGGAGGTGTTAAATAGGCTAGTG |
| attB2-Cmed092450 Rev | GGGGACCACTTTGTACAAGAAAGCTGGGTTTCACTGCGAGGCCACGGCACCCG |
| attB1-Cmed153350 For | GGGGACAAGTTTGTACAAAAAAGCAGGCTTAATGGCACTATCAAATAACTCCTTC |
| attB2-Cmed153350 Rev | GGGGACCACTTTGTACAAGAAAGCTGGGTTCTAATCAGTTTTTTTCGCTTCATT |
| attB1-Cmed153360 For | GGGGACAAGTTTGTACAAAAAAGCAGGCTTAATGGCGACAATTGAAACGCAACC |
| attB2-Cmed153360 Rev | GGGGACCACTTTGTACAAGAAAGCTGGGTTTTACTGTTCTTTGGTTATTCCATTGC |
| attB1-Cmed063180 For | GGGGACAAGTTTGTACAAAAAAGCAGGCTTAATGGGGACGCTTGTTGAGAATATC |
| attB2-Cmed063180 Rev | GGGGACCACTTTGTACAAGAAAGCTGGGTTCTACTTAGACTTATTTCTATTCTTC |
| attB1_Cmed009890 For | GGGGACAAGTTTGTACAAAAAAGCAGGCTTAATGGAGAAAGAAATAAAGCCGCGT |
| attB2_Cmed009890 Rev | GGGGACCACTTTGTACAAGAAAGCTGGGTTCTATAGAGTTTTGGTGTCATCTTC |
| attB1_Cmed022650 For | GGGGACAAGTTTGTACAAAAAAGCAGGCTTAATGTCACTCACATCAGAAGTGCC |
| attB2_Cmed022650 Rev | GGGGACCACTTTGTACAAGAAAGCTGGGTTCTATTCTTTGACCTCATCCGTGT |
| attB1_Cmed054120 For | GGGGACAAGTTTGTACAAAAAAGCAGGCTTAATGCCGCCTCTCGTGGACCCGGT |
| attB2_Cmed054120 Rev | GGGGACCACTTTGTACAAGAAAGCTGGGTTTCAATCTGCTTTCTTCTCTTCGT |
| attB1_Cmed070400 For | GGGGACAAGTTTGTACAAAAAAGCAGGCTTAATGGCTCCAAATTCAGAAACGGC |
| attB2_Cmed070400 Rev | GGGGACCACTTTGTACAAGAAAGCTGGGTTTCAGGAAGTGAATTTTTTGTACG |
| attB1_Cmed116530 For | GGGGACAAGTTTGTACAAAAAAGCAGGCTTAATGGAGACCGCCCCGGACTGCAGT |
| attB2_Cmed116530 Rev | GGGGACCACTTTGTACAAGAAAGCTGGGTTTTATTCAGTCTTGTCAGAATTGATG |
| attB1_Cmed116590 For | GGGGACAAGTTTGTACAAAAAAGCAGGCTTAATGAGTTCGTCACTACTCCTGGC |
| attB2_Cmed116590 Rev | GGGGACCACTTTGTACAAGAAAGCTGGGTTTTACTCTACTTTCTTATGGACAAT |
| attB1_Cmed116600 For | GGGGACAAGTTTGTACAAAAAAGCAGGCTTAATGGCTCCCGCCCAACAAGACGT |
| attB2_Cmed116600 Rev | GGGGACCACTTTGTACAAGAAAGCTGGGTTCTATGAATGTTTGTATTCTTCTG |
| Cmed153360-3'GSP297 | CCAAGTCACCTTTCTCCATGTCTACCAC |
| Cmed153360-5’GSP341 | GGACGCGTGGTGGTAGACATGGAG |

**Table S3 Results of normality tests for data in Figure 10a)**

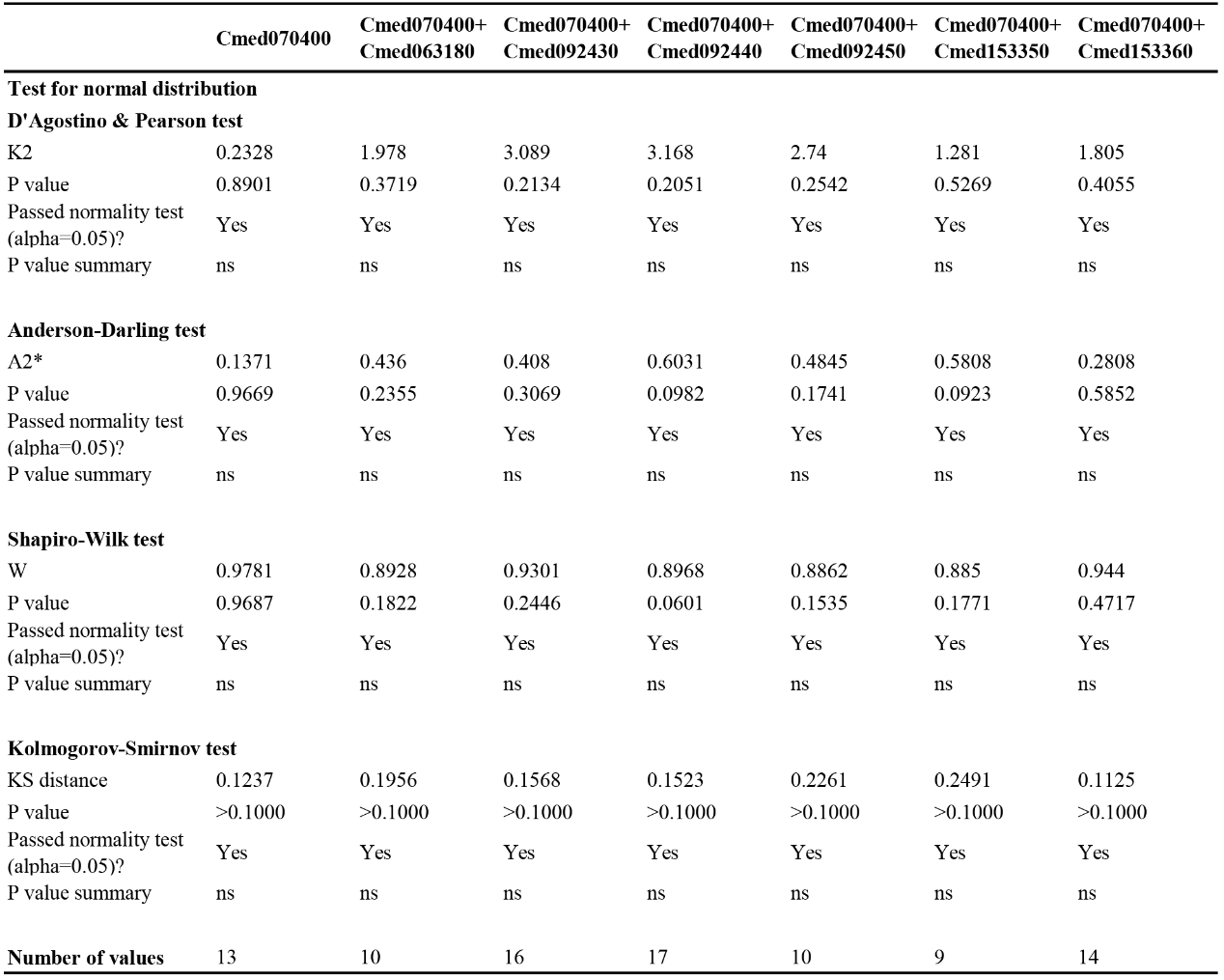

** p* < 0.05, *** p* < 0.01, ns: not significant.

**Table S4 Results of normality tests for data in Figure 10b)**

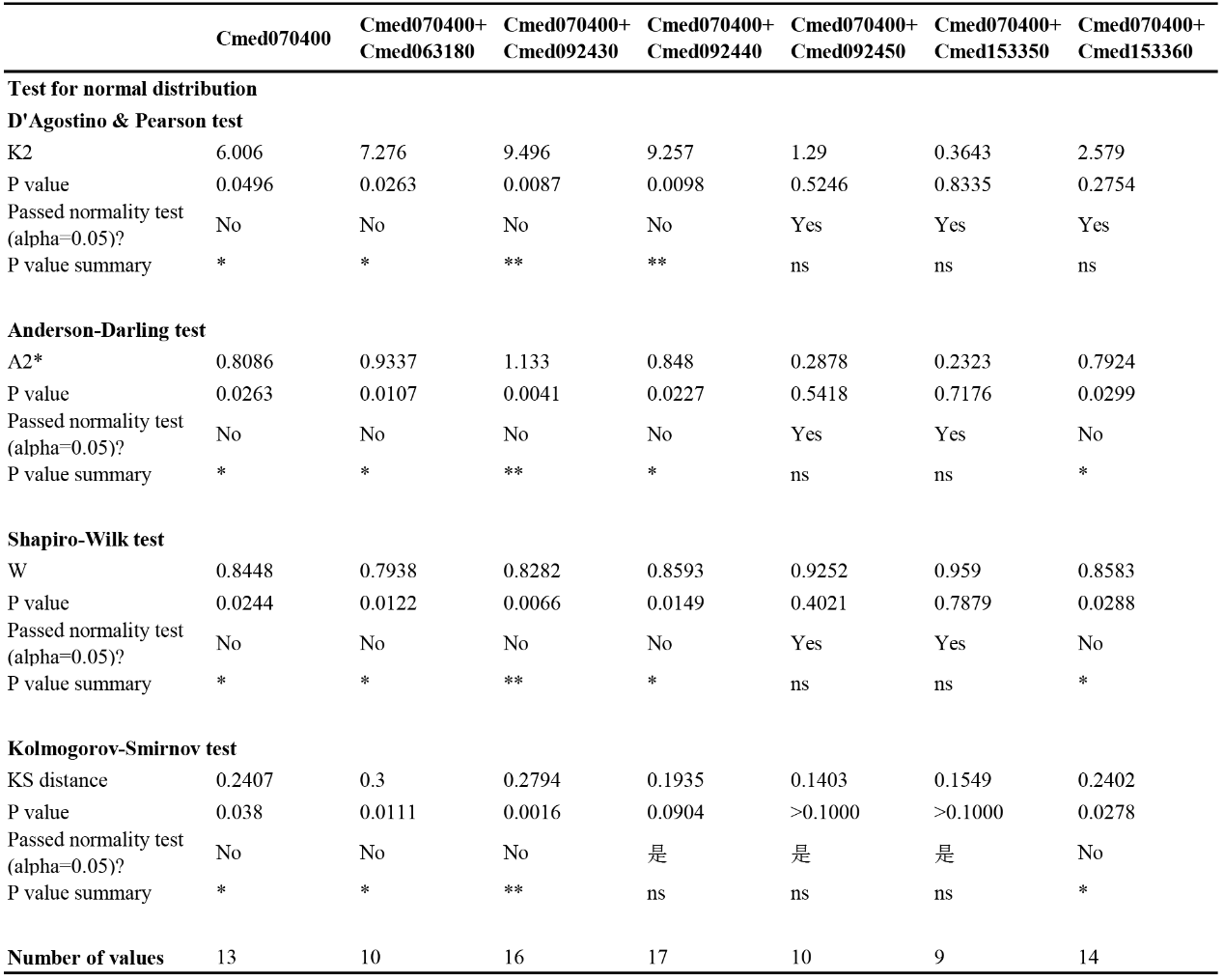

** p* < 0.05, *** p* < 0.01, ns: not significant.

**Table S5 Results of normality tests for data in Figure 10c)**

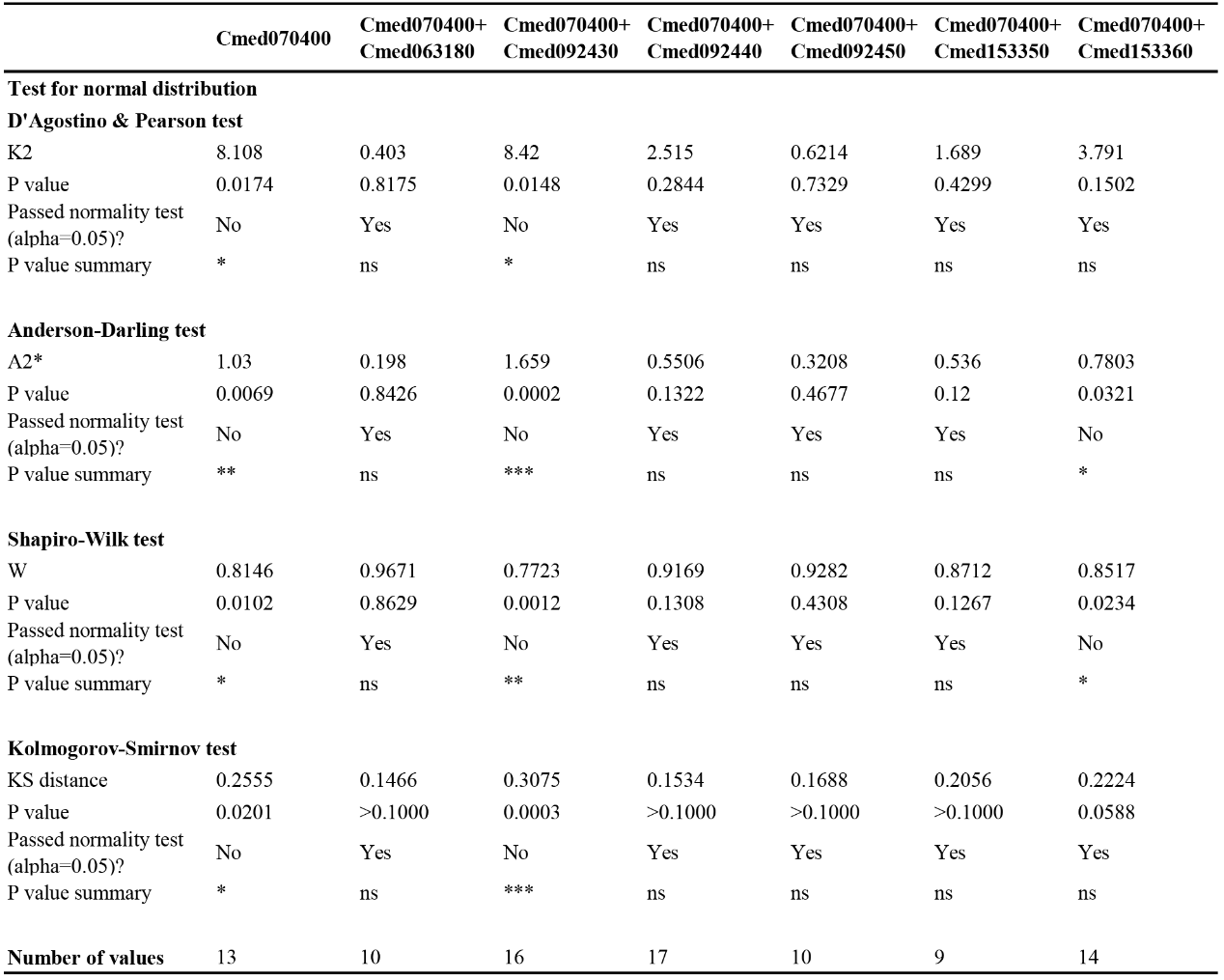

** p* < 0.05, *** p* < 0.01, ns: not significant.

**Figure S1. Heterologous expression of *IgalASE1* in *N. benthamiana*.**

Gas Chromatography-Mass Spectrometry (GC-MS) analysis of fatty acid methyl ester profiles of the leaves expressing *IgalASE1*. Native compounds from plant were shown in upright and compounds produced from the introduced elongase were shown in red. The internal standard, 19:Me, is abbreviated as IS in the figure.

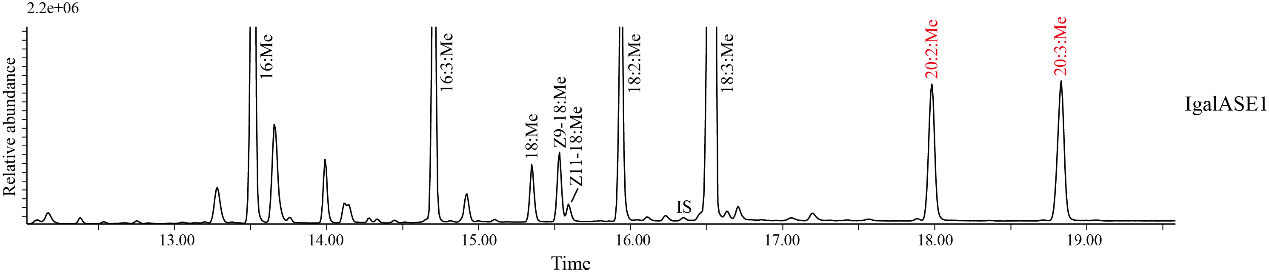

**Figure S2. Expression levels of putative pheromone biosynthesis genes in the pheromone gland of *C. medinalis* based on an independent RNA-seq dataset.** Gene expression levels are shown as Fragments Per Kilobase of transcript per Million mapped reads (FPKM). This dataset was generated from an independent RNA-seq experiment and serves as a biological replicate to validate the reproducibility of transcriptome-derived expression profiles.

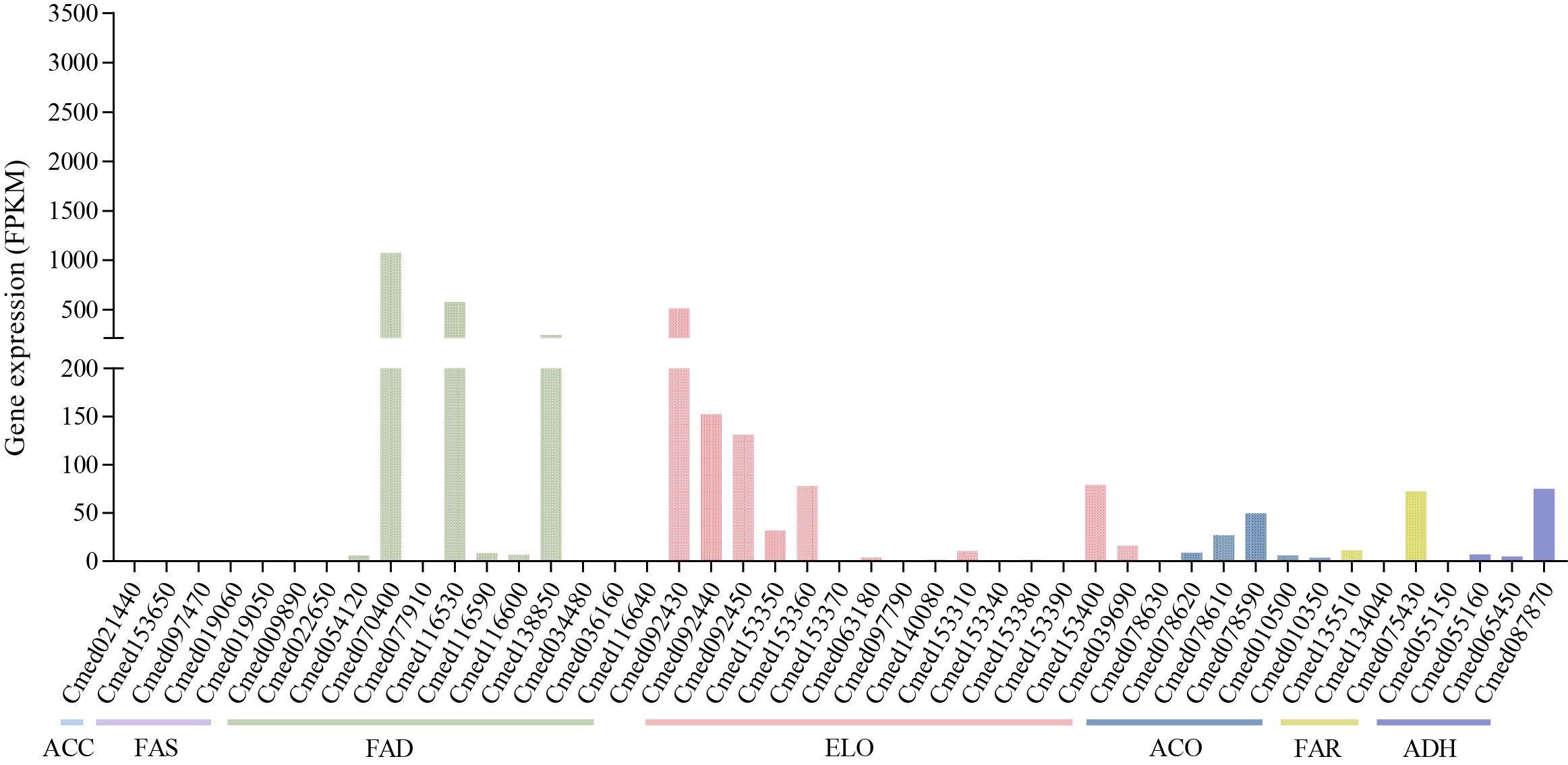

**Figure S3. Proposed pathways towards the biosynthesis of major sex pheromone component in *C. medinalis*.**

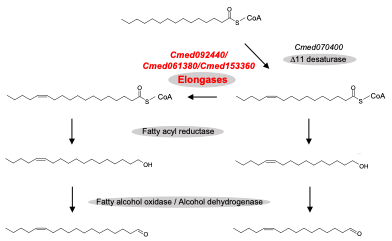
